# Graphical Representation of Landscape Heterogeneity Identification through Unsupervised Acoustic Analysis

**DOI:** 10.1101/2024.08.12.607639

**Authors:** Maria J. Guerrero, Camilo Sánchez-Giraldo, Cesar A. Uribe, Víctor M. Martínez-Arias, Claudia Isaza

## Abstract

1. Changes in land use and climate change threaten global biodiversity and ecosystems, calling for the urgent development of effective conservation strategies. Recognizing landscape heterogeneity, which refers to the variation in natural features within an area, is crucial for these strategies. While remote sensing images quantify landscape heterogeneity, they might fail to detect ecological patterns in moderately disturbed areas, particularly at minor spatial scales. This is partly because satellite imagery may not effectively capture undergrowth conditions due to its resolution constraints. In contrast, soundscape analysis, which studies environmental acoustic signals, emerges as a novel tool for understanding ecological patterns, providing reliable information on habitat conditions and landscape heterogeneity in complex environments across diverse scales and serving as a complement to remote sensing methods.
2. We propose an unsupervised approach using passive acoustic monitoring data and network inference methods to analyze acoustic heterogeneity patterns based on biophony composition. This method uses sonotypes, unique acoustic entities characterized by their specific time-frequency spaces, to establish the acoustic structure of a site through sonotype occurrences, focusing on general biophony rather than specific species and providing information on the acoustic footprint of a site. From a sonotype composition matrix, we use the Graphical Lasso method, a sparse Gaussian graphical model, to identify acoustic similarities across sites, map ecological complexity relationships through the nodes (sites) and edges (similarities), and transform acoustic data into a graphical representation of ecological interactions and landscape acoustic diversity.
3. We implemented the proposed method across 17 sites within an oil palm plantation in Santander, Colombia. The resulting inferred graphs visualize the acoustic similarities among sites, reflecting the biophony achieved by characterizing the landscape through its acoustic structures. Correlating our findings with ecological metrics like the Bray-Curtis dissimilarity index and satellite imagery indices reveals significant insights into landscape heterogeneity.
4. This unsupervised approach offers a new perspective on understanding ecological and biological interactions and advances soundscape analysis. The soundscape decomposition into sonotypes underscores the method’s advantage, offering the possibility to associate sonotypes with species and identify their contribution to the similarity proposed by the graph.

## 1. Introduction

Understanding landscape heterogeneity, which refers to the uneven distribution of elements across landscapes (Farina, 2022), is crucial when designing and evaluating the impact of conservation policies, particularly considering the increasing pressure these systems face from land use changes and global warming (Walther et al., 2002; Lambin and Meyfroidt, 2011; Powers and Jetz, 2019). This pressure has precipitated a global decline in biodiversity as essential natural habitats necessary for the survival of different species are diminished or outright eradicated (Newbold et al., 2015). Collecting biodiversity data on various regions enables an understanding of how landscape structure influences species, their behaviors, and the dynamics within heterogeneous landscapes (Worboys et al., 2010; Tscharntke et al., 2012). However, the inherent complexity of biological systems makes monitoring natural ecosystems costly and challenging, especially when the studies are conducted manually and in tropical ecosystems characterized by high biodiversity (Kallimanis et al., 2012). Therefore, a key enabling task in supporting conservation efforts is the automatic identification of the intricate interactions within landscapes to estimate their heterogeneity, thereby supporting conservation efforts.

Landscape heterogeneity identification has been approached from a spatial perspective using coverage and spectral indices obtained from satellite imagery (Ndao et al., 2021; Radocaj et al., 2020). These spectral indices enable monitoring vegetation properties by identifying specific land cover types or their distinct properties, as reflected in unique spectral values compared to the surrounding area (Radocaj et al., 2020). However, the scale these indices operate is unsuitable when the required analysis goes beyond vegetation and coverage types. Moreover, pixel-based analysis from satellite imagery might lead to misclassifying vegetation cover, especially considering its conservation status, and is ineffective at characterizing undergrowth conditions due to inherent scale limitations(Montero et al., 2023; Berra and Gaulton, 2021; Al-Wassai and Kalyankar, 2013).

Using a different modality, soundscape analysis has become one of the more common alternatives to measuring ecological processes. This approach is based on studying how sounds from various sources, such as biological organisms, geophysical phenomena, and human activities, can be used to understand these processes at different temporal and spatial scales (Fuller et al., 2015; Pijanowski et al., 2011). In this context, Passive Acoustic Monitoring (PAM) has become popular in land environments, employing sound to monitor wildlife and enabling an understanding of their dynamics (Dumyahn and Pijanowski, 2011; Sueur and Farina, 2015; Gibb et al., 2018; Sugai et al., 2019; Stowell and Sueur, 2020). This type of monitoring has been made possible through advancements in digital recording technology, enabling the remote, autonomous, and effective collection of acoustic activity in studied ecosystems (Acevedo and Villanueva-Rivera, 2006; Sousa-Lima et al., 2013). Additionally, massive datasets are generated due to their capacity for deployment in extensive areas over prolonged periods, offering a broad and detailed perspective on changes and patterns in the soundscape.

PAM data have generally been used to monitor sound-emitting species through automatic detection methodologies using machine learning and deep learning techniques (Bedoya et al., 2014; Zhao et al., 2017; LeBien et al., 2020; Dufourq et al., 2020; Nolasco et al., 2023). Regarding the analysis of the soundscape and its relationship with ecological processes, different methodologies have been proposed to discriminate soundscape components (Bellisario et al., 2019), identify various types of habitats or coverages (Gómez et al., 2018; Apoux et al., 2023; Castro-Ospina et al., 2024), and differentiate between the transitional states of ecosystems Rendon et al. (2022); Castro-Ospina et al. (2023). These proposals have in common the use of supervised learning methods, requiring labels of the habitat type or the state of transformation for the learning process. Relying on labels for the learning process in biological and ecological applications represents a problem because there is a high degree of uncertainty in the phenomena, and the states of the ecosystems are not necessarily known a priori. Limiting the analysis to identifying specific habitat types or discrete changes in ecosystems restricts the understanding of the phenomenon (Rendon et al., 2022).

Several acoustic indices have been developed to represent the complexity of the landscapes based on their features, such as species richness, sound dissimilarities between remote areas, and anthropogenic activity, among others (Gasc et al., 2015). These indices have been used to identify dissimilarity among types of coverages (Barbaro et al., 2022; Hayashi et al., 2020), provide information about habitat changes (Sánchez-Giraldo et al., 2021), and analyze the behaviors of acoustic communities (Pijanowski et al., 2011), by analyzing the relations between acoustic diversity generated by the indices and the heterogeneity in composition and configuration of the landscape in different habitats. However, despite their utility in evaluating the acoustic properties of the landscape without requiring labels, their applicability as a proxy for acoustic biodiversity has been recently questioned, as contradictory patterns have been observed in different geographical regions and environments (Llusia, 2024; Bicudo et al., 2023; Alcocer et al., 2022).

Previous studies (Burivalova et al., 2019) explore how industrial logging affects animal vocalization diversity in tropical forests using soundscape recordings to assess alpha and beta diversity through soundscape saturation and dissimilarity measures based on acoustic indices. They note the lack of a standardized sound similarity measure, as discussed in (Bormpoudakis et al., 2013), and suggest that higher similarity in recordings from areas of low diversity implies subtractive homogenization. Conversely, lower similarity paired with high diversity indicates additive heterogenization. On the other hand, (Bormpoudakis et al., 2013) demonstrated that ambient sound is not only spatially heterogeneous but also directly related to habitat-type structure. They provided evidence of habitat-specific acoustic signatures using unsupervised learning techniques, showing that each habitat type has a unique soundscape that can be used to identify and monitor ecological processes. This approach underscores the potential of using acoustic properties to define and differentiate habitat types beyond traditional species-specific concepts. Based on these findings, no previous research has identified landscape heterogeneity by analyzing biophony by decomposing the landscape into acoustic entities without relying on species-specific approaches, acoustic indices, or general landscape measures.

In this work, we aim to address the challenge of estimating heterogeneity among different geographical locations using unlabeled data from passive acoustic monitoring. By building on the structural properties of the data, we leverage network science techniques, which have been employed to understand complex systems such as the human brain, where some analysis identifies similarities between different brain regions (as in The Human Connectome Project^1^), genomic analysis, disease evolution, ecological networks, and social networks (Li et al., 2021; Gao et al., 2021; Fu et al., 2021; Zhou et al., 2022). Our approach uses a sparse Gaussian graphical model, Graphical Lasso (GLasso) (Friedman et al., 2008), which generates a sparse precision matrix containing similarity information. GLasso has been effectively applied in various fields, including psychology, gene analysis, and neuroscience (Bhushan et al., 2019; Huang et al., 2020; Ranciati et al., 2021), where it handles different types of input data. This method is advantageous because it forces small partial correlation coefficients to zero, inducing sparsity in the graph and making it easier to interpret by reducing the number of links (Friedman et al., 2008). This method allows for the establishment of similarity between variables and, through a graph representation, illustrates the heterogeneity or similarity between different geographical locations, where nodes represent a distinct geographical location and links indicate the presence of a relationship between sites. The key novelty of our analysis lies in decomposing the soundscape into acoustic entities called sonotypes, which are sound patterns occupying the same time-frequency acoustic space, using the unsupervised methodology by (Guerrero et al., 2023). It showed that identifying sonotypes at a location corresponds to their biophony and changes according to the time of day. By counting the occurrences of each sonotype at different sites, we can determine the acoustic structure of each location. This structure is a unique acoustic fingerprint, encompassing all frequency bands and enabling a comprehensive sound-scape analysis to describe the landscape’s sound. Each site is characterized by the variety of sonotypes present and their proportion throughout the day relative to others. This pattern creates a unique signature for each location, allowing comparisons between different sites. Using these acoustic structures as input features, we leverage network science techniques to identify and visualize the relationships between geographical sites. We analyze the relationships based on the sonotypes and their temporal and frequency distribution.

We validate the proposed approach in two case studies. In the first case, we apply the graph model to acoustic structures extracted from an acoustic dataset acquired through passive acoustic monitoring in an agricultural region in Colombia. The second case study involves analyzing the same acoustic dataset but at different times of the day to identify changes in the graph according to soundscape changes during the day. Correlating our findings with ecological metrics such as Bray-Curtis dissimilarity estimation (Bray and Curtis, 1957; Deichmann et al., 2017) and spectral indices obtained from satellite imagery (Radocaj et al., 2020) at the studied site reveals significant insights into landscape heterogeneity. Our approach illustrates the contribution of sonotypes to the similarity patterns identified by the graph. It shows that acoustic monitoring, enhanced by network science techniques, complements traditional ecological methods, offering a nuanced understanding of biodiversity and ecosystem dynamics.

## 2. Materials and Methods

### 2.1. Unsupervised acoustic heterogeneity identification and graphic representation

The analysis presented in this study encompasses a series of steps (Figure 1) aimed at identifying heterogeneity among geographical sites using passive acoustic monitoring data. First, a signal analysis was carried out to identify and remove sound files predominantly associated with geophony, specifically intense rain events, using the rain detection algorithm proposed by (Bedoya et al., 2017). Subsequently, we applied the unsupervised methodology proposed by (Guerrero et al., 2023), a comprehensive acoustic analysis framework involving an initial preprocessing step designed to reduce background noise, followed by the automatic decomposition of recordings into sonotypes, acoustic entities composed of unique frequencies and time frames. Using these identified sonotypes, we constructed an acoustic structure for each site, represented as an *m* × *n* matrix indicating the number of occurrences of each sonotype (*n*) at each site (*m*). Afterward, acoustic similarities among geographical locations were identified using a graph inference technique (Graphical Lasso - GLasso) based on the acoustic structure. Finally, we performed an analysis seeking to interpret the links created by the model from sonotypes and from an ecology perspective.

**Figure 1.**
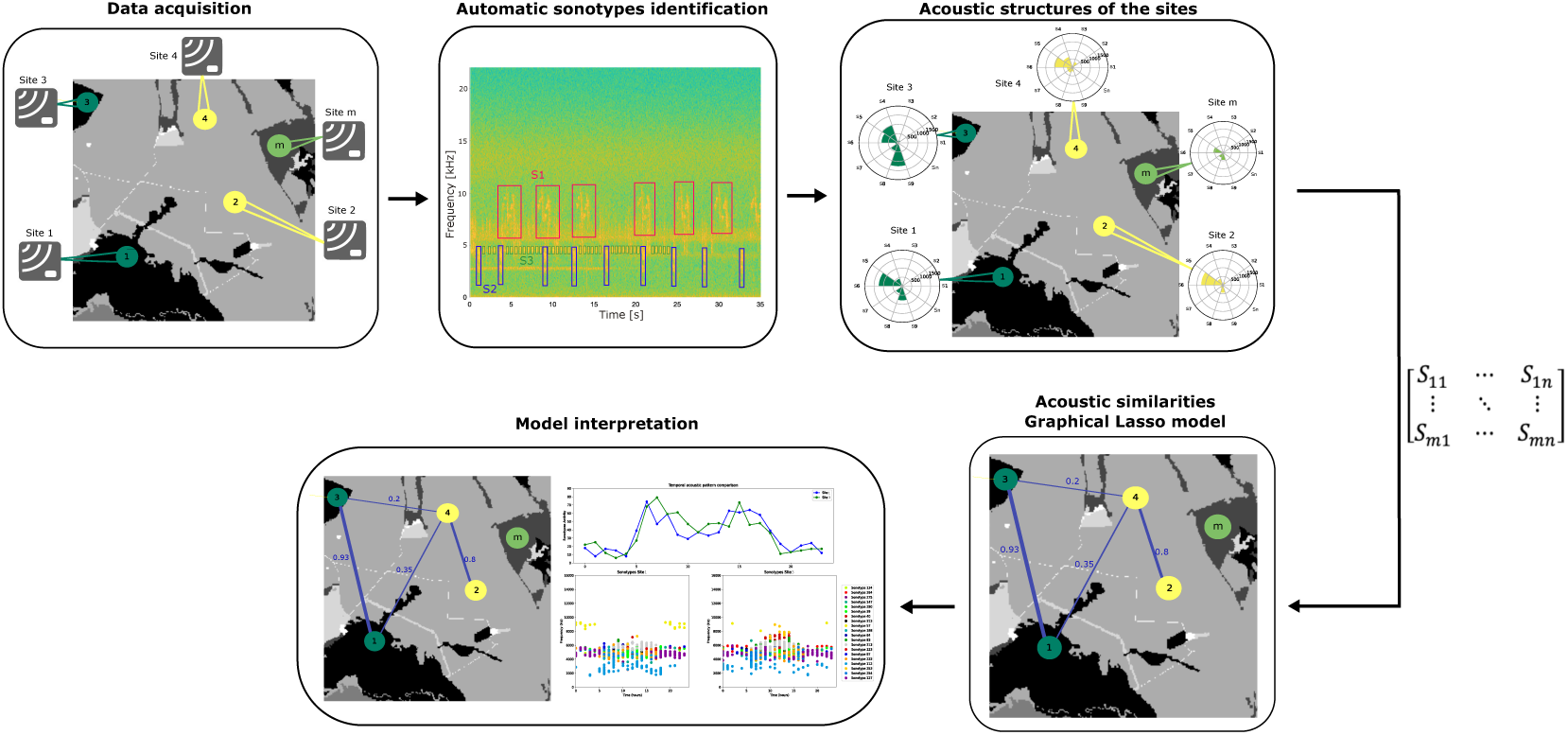
Unsupervised acoustic heterogeneity identification framework. It begins with the input data, composed of acoustic data from different geographical sites. This is followed by segregating this data into unique acoustic entities, or sonotypes. These sonotypes are then used to map the acoustic structure of each location, forming a matrix that quantifies the presence of each sonotype per site. The process continues with a Gaussian graphical model to discern acoustic similarities between the sites, culminating in an analysis that interprets these relationships from an ecological perspective.

We developed an evaluation case by simulating a soundscape to analyze the generality of the graph model and establish a ground truth. In this case, we generated various acoustic structures by creating sonotypes and their occurrences in hypothetical locations, examining scenarios where sites were either expected or not expected to be connected in the model (e.g., similar acoustic structures between sites, different acoustic structures, the same sonotypes but with varying numbers of occurrences, etc.). The design and evaluation of this evaluation case are presented in Appendix A, demonstrating the correct functionality of the graph model used in this study.

Additionally, we tested the performance of the graph methodology in two case studies. The first case study applied the proposed framework to the acoustic data described in Section 2.2.1 (Puerto Wilches dataset). Then, to analyze the temporal influence of the Puerto Wilches soundscape, we examined it by time slots, forming case study 2 (Section 2.2.2). Finally, to validate the performance of our proposed methodology, we conducted a heterogeneity analysis by comparing our graph inference results with other commonly used methods. This comparison involved aligning our findings with spectral indices derived from satellite imagery, performing a Bray-Curtis dissimilarity analysis, and calculating the Spearman correlation coefficient.

#### 2.1.1. Automatic sonotypes identification

The input features for the network inference model are the acoustic structures of each sampled site. This acoustic structure is determined by the sonotypes found at each site and their number of occurrences.

To estimate the sonotypes present in the soundscape (acoustic data), we use (Guerrero et al., 2023) proposal. This unsupervised methodology automatically segments the acoustic activity present in the audio recordings and clusters them based on their acoustic features (frequency information and cepstral coefficients) similarities. The resulting clusters exhibit distinct acoustic patterns that can be associated with species calls, referred to as sonotypes. Additionally, in (Guerrero et al., 2023), it was demonstrated that it is possible to characterize biophony through sonotypes similarly to how acoustic indices such as ACI, BI, NP, and SO indices do.

In this approach, sonotypes are not directly linked to specific species calls. Instead, we focus on working with sonotypes as descriptors of the soundscape and utilizing their occurrence frequencies to generate the acoustic structure of each site. This methodology does not require any parameterization.

#### 2.1.2. Acoustic structures of the sites

Once sonotypes were obtained, we generated the acoustic structures of each study site by compiling the number of occurrences of each sonotype (*n*) in each sampled site (*m*), thus resulting in a matrix *m* × *n*. Each row of the matrix corresponds to a unique site (Figure 2). This graphical representation illustrates the occurrence of identified sonotypes at a particular site. Each bar corresponds to a unique sonotype (*S* 1*, S* 2*, …, S n*) obtained automatically from (Guerrero et al., 2023) proposal. The length of each bar is proportional to the number of occurrences of sonotypes, providing a rapid view of the predominant acoustic patterns at the site.

**Figure 2.**
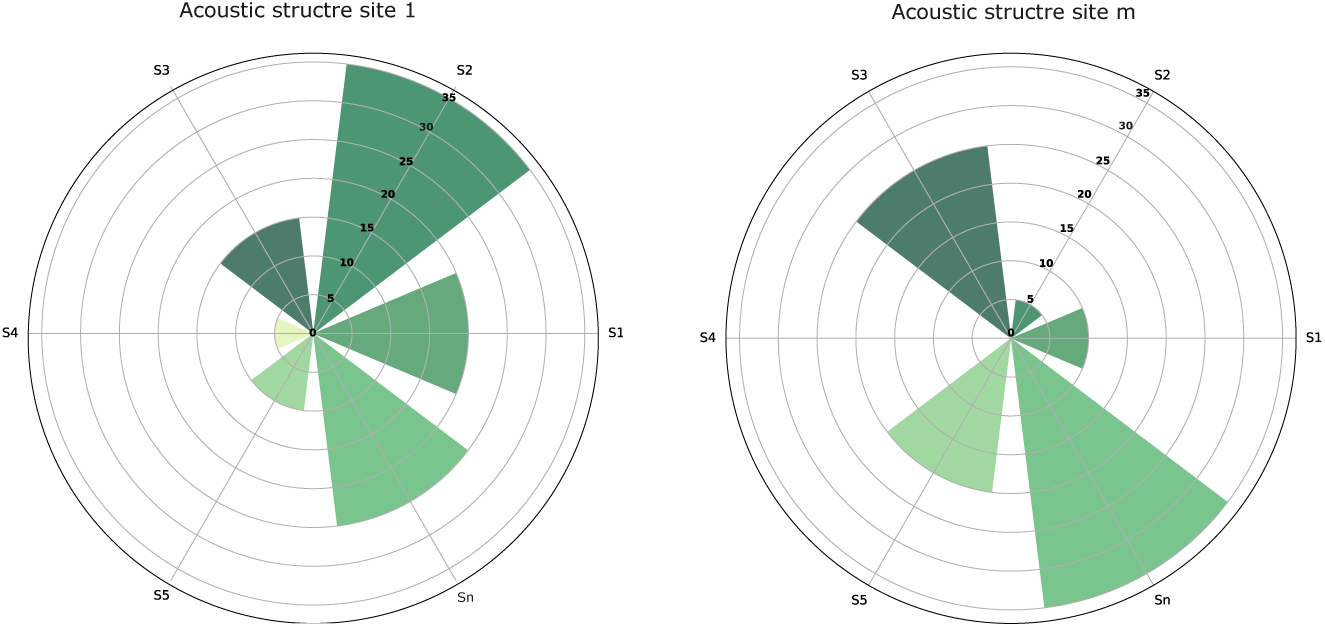
Example of the acoustic structures. Figure (a) illustrates the acoustic structure of site one, while figure (b) displays a similar structure from site m. Each radial bar, denoted as *S* 1 to *S n*, corresponds to a unique sonotype detected in the soundscape, and it is represented in different colors, with its length indicating the number of occurrences. The chart serves as an acoustic fingerprint, highlighting the diversity of sonotypes and providing a visual review of the biophony of the site. The variation in sonotype occurrences across the two sites enables comparative analysis, showcasing that even sites with similar acoustic entities may exhibit different sonotype occurrence patterns.

These acoustic structures facilitate both quantitative comparisons of sonotypes within the same site and across different sites. By examining these structures, experts can identify the richness of sonotypes, acoustic diversity, and potential differences in biophony composition among various geographical sites. This method offers comprehensive insights into the acoustic behavior of sonotypes at each site. It provides a nuanced understanding of the soundscape, enhancing our ability to interpret and compare the complex acoustic environments of natural habitats.

#### 2.1.3. Gaussian graphical model for acoustic heterogeneity identification

Gaussian graphical models (GGM) comprise several features or variables represented by nodes and links showing relationships among those features or variables. Thus, the absence of a link shows a nonexistent relationship among variables.

Graphical Lasso estimates a precision matrix by applying *L1* (Lasso) regularization to the elements. To estimate the precision matrix Θ, GLasso assumes multivariate Gaussian distribution with mean *µ* and covariance matrix Σ (Friedman et al., 2008).

Consider a scaled and centered data matrix **X** ∈ R*^m^*^×*n*^ where *n* measures the occurrence of the sonotypes in each sampled site *m*, and Σ ∈ R*^m^*^×*m*^ its covariance matrix. The algorithm aims to estimate a sparse precision matrix Θ = Σ^−1^ where Θ ∈ R*^m^*^×*m*^ is the inverse covariance matrix and correspond to pairs of variables that are conditionally independent. The conditional dependence relationships can be represented by a graph where nodes represent the sites (in the case of our study), and edges connect a pair of nodes based on their relationship (Mazumder and Hastie, 2011). To calculate the precision matrix, GLasso problem minimizes a *L1*-regularized negative log-likelihood as is showing in Eq.(1):

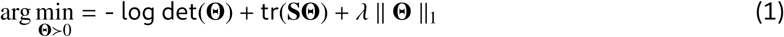

Where **S** is the sample covariance matrix calculated from sonotypes matrix **X**, ∥̄ Θ ∥̄_1_ represents the sum of the absolute values of Θ, *λ* is a tuning parameter which control the sparsity of Θ.

To estimate the precision matrix using Graphical Lasso, we used the ‘GraphicalLassoCV’ function from the ‘scikit-learn’ library version 1.5.1 in Python (Pedregosa et al., 2011). This function performs tuning parameter selection for *λ* through cross-validation. Once the precision matrix has been obtained, we used ‘NetworkX’ library version 3.2.1 in Python (Hagberg et al., 2008) to create the graph that shows the relationship between geographical locations based on Θ results.

Due to the sparsity induced by GLasso, many values in the precision matrix are zero. However, a threshold is defined to determine the links that remain. Only the values that exceed this threshold are considered, ensuring that statistically significant correlations are retained, reducing the likelihood of spurious connections. Empirically, we validated the threshold by comparing the resulting network structures against known ecological relationships, ensuring that the selected threshold provides a meaningful and interpretable representation of the data.

#### 2.1.4. Acoustic heterogeneity identification analysis

Once we identify the similarities among sites due to the links created by the Graph Lasso model, we interpret the model using the sonotypes that constitute the acoustic structure and its information. Given the significant number of sonotypes, we analyzed matrix **X** to estimate the Total Variation Distance (TVD) to identify the most relevant sonotypes in each pair of connected sites. TVD measures the difference between two probability distributions and is presented as follows:

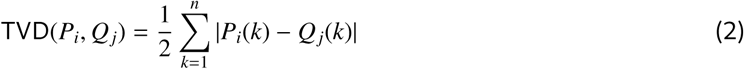

Where *P_i_*and *Q_j_* represent the relative frequencies of sonotype *k* at sites *i* and *j*, respectively. Here, *n* denotes the total number of sonotypes considered in the analysis, consistent with the value obtained from section 2.1.2. For each pair of sites *i* and *j*, the counts of detected sonotypes are extracted, and their relative frequencies are calculated by dividing each sonotype counting by the total count of all sonotypes at that site. A TVD value of 0 indicates that the two distributions are identical (i.e., the relative frequency or proportion of each sonotype is the same in both sites). In contrast, 1 indicates the maximum possible difference between them. The result is a new matrix **D** ∈ R*^p^*^×*n*^ with the TVD values of each sonotype *n* at each relevant site pair *p*.

To identify the most representative sonotypes from matrix **D**, we first calculate the median and standard deviation of the TVD for each sonotype across each pair of sites. We then define a threshold for selecting relevant sonotypes by adjusting the median downward by 0.02 times the standard deviation. This specific threshold was chosen based on both theoretical considerations and empirical validation. Theoretically, it ensures that sonotypes with low variability are prioritized, which are likely to be more representative of the acoustic structure of the sites. Sonotypes with TVD values below this threshold are considered significant.

Having identified the most representative sonotypes for each pair of sites, we utilize their frequency-time information (from Section 2.1.1) to create a visual representation that will help to understand the link between a pair of sites. This was achieved in two complementary ways: We plotted individual sonotypes as colored dots and crosses on a scatter plot, with the position of each dot corresponding to the hour of detection and its peak frequency. Dots represent sonotypes significantly related to similarity, while crosses denote sonotypes associated considerably with differences among pairs of sites. Alongside this, by aggregating the count of sonotypes detected at each hour throughout the day, we produced a line graph that depicts the temporal acoustic pattern for each site. This representation allows an understanding of the GLasso links by identifying common sonotypes at the sites and their distribution in the soundscape.

The algorithms implemented for this proposal are available at: Approach algorithms here.

### 2.2. Case studies

We tested the capabilities of the Graphical Lasso model to estimate similarities from acoustic structures among different geographical locations using two case studies. In case study 1, we analyzed data from passive acoustic monitoring related to an oil palm plantation site. In contrast, in case study 2, we examined the same acoustic data but in different time slots to assess their temporal influence.

#### 2.2.1. Case 1: Puerto Wilches acoustic dataset

In this case study, we used a real acoustic database derived from passive acoustic monitoring conducted in a rural area of the municipality of Puerto Wilches, Santander, Colombia (7^◦^21^′^52.5”*N,* 73^◦^51^′^33.0”*W*). The landscape of this area is predominantly covered by oil palm plantations of different ages (75%). It also features a mix of secondary vegetation (7.6%), patches of forest (6.13%), grasslands (5.5%), and areas of aquatic vegetation (3.2%). Although the region includes several buildings and a network of secondary roads serving oil palm plantations and livestock farming, anthropogenic components are notably low due to minimal human activity. These roads are private, with infrequent use primarily limited to plantation personnel. Residential buildings are sparse and distant from recording locations, reducing the likelihood of significant anthropogenic sound disturbances.

The dataset consists of 19,598 audio recordings collected over 10 days in March 2021 (dry season) from 17 sites within the selected area (see Figure 3). A Song Meter Mini device (Wildlife Acoustics, Inc.) was used for data collection at each sampling site, programmed to record one minute every ten minutes with a sampling rate of 48 kHz. Each recorder was placed at a minimum distance of 300 m from the other. Considering the 9 km^2^ extent of the study area, this spacing aimed to ensure spatial independence among recordings by minimizing the influence of shared sound sources and obtaining a representative sample of the acoustic variability across the landscape. In Figure 3A, the general location and the different types of cover of the sites are shown, and in Figure 3B, the arrangement and geographical location of the recorders used are displayed.

**Figure 3.**
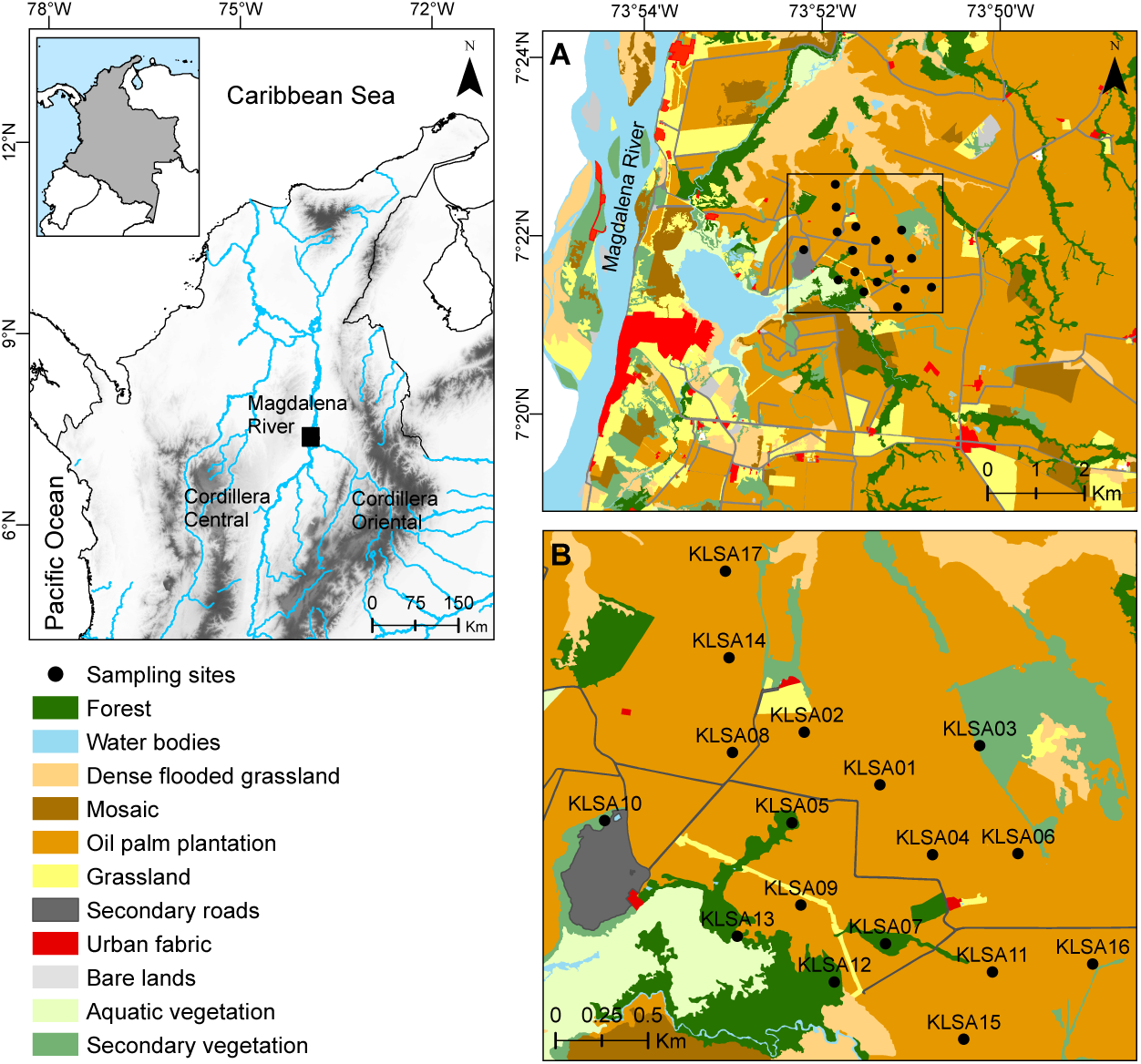
Study site in Puerto Wilches, Santander, Colombia. Spatial distribution of the recorders and landscape cover type. Each white dot represents a recorder with its given name

In this case study, we address the problem of identifying similarities among sites using acoustic data in an unsupervised way. In this context, we do not have a ground truth graph that represents expert-defined connections. Instead, we have the acoustic dataset and the classification of coverages where the data were collected.

This dataset was previously analyzed in Guerrero et al. (2023), where the identified sonotypes exhibited trends similar to acoustic indices commonly associated with biophony. These findings support the assumption that, given the characteristics of the study area, the acoustic patterns extracted through sonotypes predominantly reflect biophonic activity.

#### 2.2.2. Case 2: Puerto Wilches temporal analysis

From an acoustic perspective, landscapes include diverse sounds arising from biophonic (sounds produced by animals), geophonic (environmental sounds such as rain or wind), and anthropophonic (human-related) sources. Although all these components can coexist, biophony clearly dominates the acoustic environment in our specific study area (see Section 2.2.1). Given this biophonic predominance, the acoustic landscape exhibits inherent daily dynamics, reflecting variations in species activity and vocalization patterns throughout the day.

Previous analyses conducted in other ecosystems have shown that dividing acoustic communities into distinct temporal segments provides a better approximation for understanding their intrinsic variability and daily community dynamics (Deichmann et al., 2017; Sá nchez-Giraldo et al., 2021; Rendon et al., 2022; Barbaro et al., 2022). Considering these precedents, our second case study aims to dissect the acoustic structure of the sites across four distinct time frames: dawn (5-8), day (8-16), dusk (16-20), and night (20-5). The sonotypes found in case 1 (2.2.1) were used in this case. Here, the counting of occurrences at each site is carried out, taking into account each time slot previously established.

To analyze this scenario alongside case study 1, we calculate the graph density, defined as the proportion of existing links relative to the total number of possible links. This metric provides insights into how connectivity patterns among sites change throughout the day compared to the overall connectivity patterns obtained when analyzing data across all daily periods.

The acoustic dataset used for case studies 1 and 2 is available at: (Datasets here.

### 2.3. Acoustic heterogeneity comparison with other methods

To perform a comparison with the Graphical Lasso method (applied to acoustic structures), we generated a graph representation from two different methods commonly used in ecology to measure similarities and estimate the heterogeneity of the landscape (Legendre and Legendre, 1998). The first method used is called Bray-Curtis dissimilarity (Bray and Curtis, 1957) using acoustic structures derived from evaluation case presented in Appendix A and case study 1, and the second method is the remotely sensed indicators or spectral indices (Jinru and Su, 2017; Radocaj et al., 2020) associated with site evaluated in case study 1.

#### 2.3.1. Comparison method using acoustic structures and Bray-Curtis dissimilarity

This method is considered reliable for quantifying differences between ecological abundance data collected from multiple sampling locations. The computation of this Bray-Curtis dissimilarity method involves summing the absolute differences between the counts and dividing this by the sum of the abundances in two sites. The formula for calculating Bray-Curtis dissimilarity **d** between site *i* and *i’* is presented in Eq.(3) where the counts are denoted by **n***_i j_* and their sample totals are **n***_i_*_+_.In this case, we used the acoustic structure matrix described in section 2.1.2.

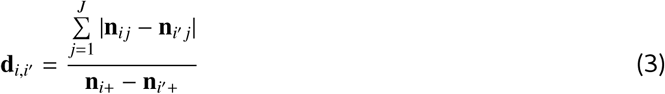

When sites are identical: **n***_i j_* = **n***_i_*_′_ *_j_*, the dissimilarity value will be 0, and 1 otherwise.

To estimate the dissimilarity matrix for all sites, the normalized acoustic structures matrix will be used as input data, and it will be computed using the **pdist** function from SciPy library version 1.13.1 in Python (Virtanen et al., 2020).

#### 2.3.2. Comparison method using remotely sensed indicators or spectral indices

These indices are obtained from remote sensing as satellites by capturing electromagnetic wave reflectance information from canopies. They are used to perform evaluations of vegetation cover, growth dynamics, conservation, and monitoring ecosystem health, among others (Jinru and Su, 2017). Table 1 presents the vegetation indices used to estimate similarities between the different geographical sites.

**Table 1:** Spectral indices generated by satellite imagery used to identify similarities between geographical locations.

| Index | Definition | Reference |
| --- | --- | --- |
| AD | Canopy Height | (Potapov et al., 2021) |
| NDRE | Normalized Difference Red Edge | (Gitelson and Merzlyak, 1997) |
| EVI | Enhanced Vegetation Index | (Huete et al., 2002) |
| COB | Coverage | (Fitzgerald et al., 2010) |
| SLAVI | Specific Leaf Area Vegetation Index | (Ali et al., 2017) |
| NDMI | Normalized Difference Moisture Index | (Gao, 1996) |
| SL | Slope | (Equator Studios, Accessed: 2023) |
| BR (MSBI) | Misra Soil Brightness Index | (Xue and Su, 2017) |
| NDBI | Normalized Difference Built-up Index | (Zha et al., 2003) |
| NDVI | Normalized Difference Vegetation Index | (Kriegler et al., 1969) |
| DI | More distance grass area greater connectivity | (Gao, 1996) |
| DN | Distance forest patches using COB | (Gao, 1996) |
| ICHN | Human Footprint Index | Correa Ayram et al. (2019) |
| FM | Fragmentation | (Jaeger, 2000) |

These indices were calculated on the Google Earth engine platform using Sentinel 2 satellite with a 10×10 pixel resolution. Subsequently, in QGIS software, a 100m buffer was applied to each sampled site, and the median value of each index was computed for all locations. This process resulted in a 17 × 14 matrix (17 locations and 14 indices). The normalized version of this matrix will be the input data for Graphical Lasso analysis to estimate the comparisons.

## 3. Results

### 3.1. Case 1: Puerto Wilches dataset

Once the performance of the method was proven using simulated data with ground truth (see Appendix A, we evaluated Graphical Lasso performance on a real ecoacoustic dataset. Here, we analyze the connections between the sites based on the sonotypes present at each connected site (or unconnected). As an unsupervised case, we do not know the expected connections a priori; for this reason, we interpreted the connected sites through their sonotypes and sonotypes time-frequency information, and then we compared the results with the other two methods commonly used in ecology to identify similarities among geographical zones.

#### 3.1.1. Acoustic structures

The acoustic dataset was analyzed following the descriptions presented in Sections 2.1.1 and 2.1.2. As a result, we obtained a 17 × 292 matrix that describes the number of occurrences of the 292 sonotypes present in each of the 17 sites during the 10 days of recording. Thus, each place will be described by its acoustic structure, as in the example presented in Figure 4, where only the first ten sonotypes (the same ten for each place) were taken to a better representation. The sonotype occurrence matrix was normalized and used as input data for Graphical Lasso and Bray-Curtis comparison analyses.

**Figure 4.**
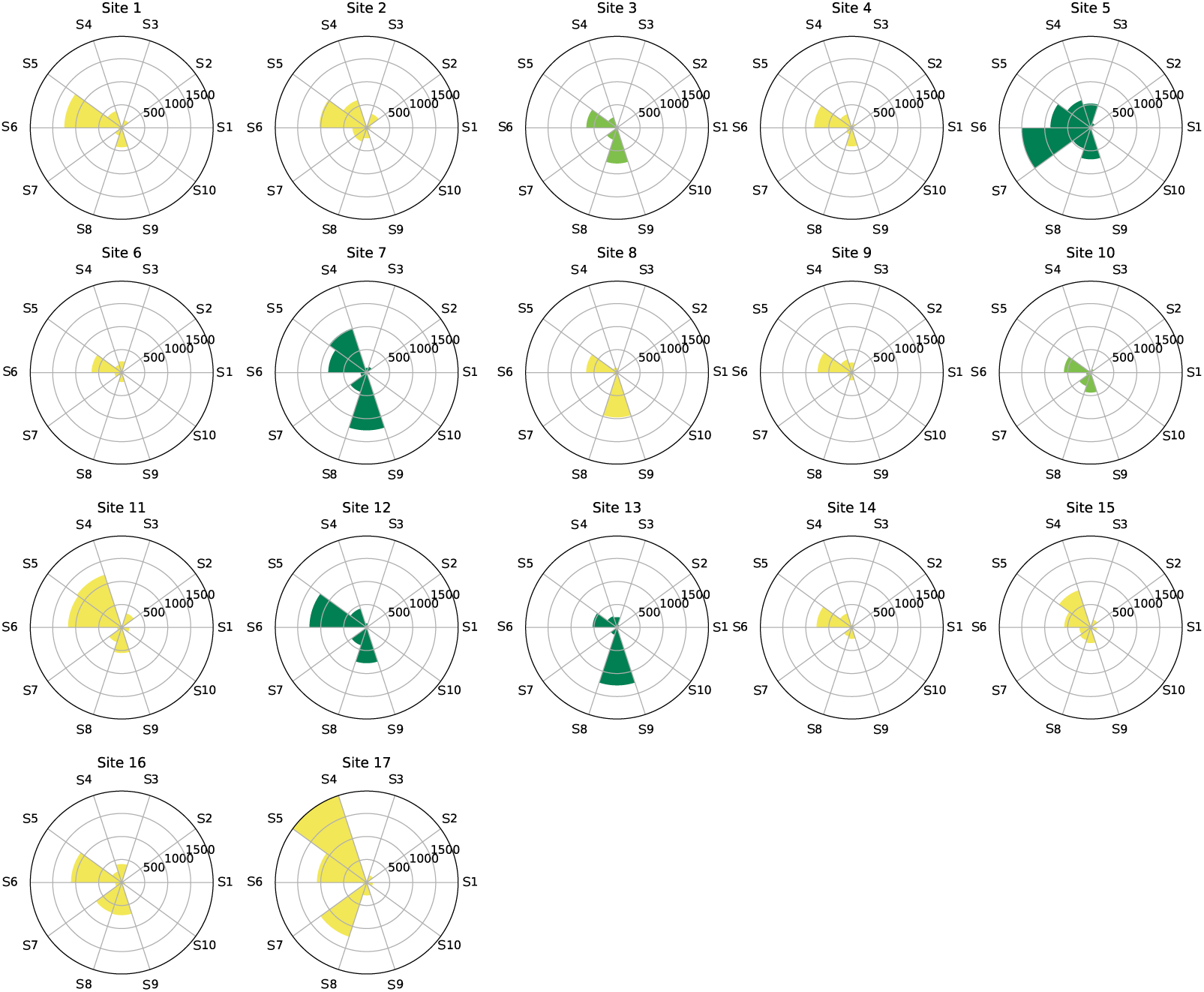
For each sampled site in Puerto Wilches, Santander, Colombia, acoustic structures were created, each consisting of the same sonotypes. As determined by expert documentation, colors across these structures represent the land cover type, where yellow represents oil palm plantations, light green represents secondary vegetation, and dark green represents forests. The coverage type information was not used to estimate the acoustic heterogeneity or connections among sites.

#### 3.1.2. Graphical Lasso model

Utilizing this network inference method, we generate a 17 × 17 precision matrix that provides information about the similarities among the analyzed locations. The Graphical Lasso model optimizes the estimation of the precision matrix by penalizing the *ℓ*_1_-norm of its elements, controlled by a regularization parameter *λ*. In our implementation using scikit-learn’s GraphicalLassoCV, this regularization parameter is named alpha. The optimal alpha found through cross-validation was 0.025, corresponding to *λ* = 0.025 in the mathematical formulation. We considered only absolute precision matrix values above 0.3 to construct the final network graph. Values above this threshold indicate strong partial correlations, highlighting the most robust connections among sites and ensuring a sparse, interpretable representation of the landscape structure.

In this case, considering that we have information regarding each place’s geographical coordinates and land cover classification, we use this data to represent the nodes in the network. Each node is positioned according to its corresponding geographical location (see Figure 3), where each color indicates its land cover type. Specifically, yellow represents oil palm plantations, light green represents secondary vegetation, and dark green represents forests. The width of the edges in the network corresponds to the higher values of the precision matrix, denoting strong connections between the two sites.

Figure 5 depicts the Graphical Lasso model using acoustic structure information, illustrating acoustic similarities across different geographical sites. This model highlights the uniformity within the oil palm plantation sites (dark gray sites in the Figure) and reveals a link between two of the four forest areas (black sites in the Figure). Notably, the distinct nodes 4 and 6 are associated with a smaller forest segment, potentially affected by the edge effect and the nearby presence of oil palm plantations or other land cover types. On the other side, nodes 2 and 8 are strongly connected, showing sonotypes in common and similar acoustic activity despite the land cover difference and geographical distance at which they are located (see section 3.1.5).

**Figure 5.**
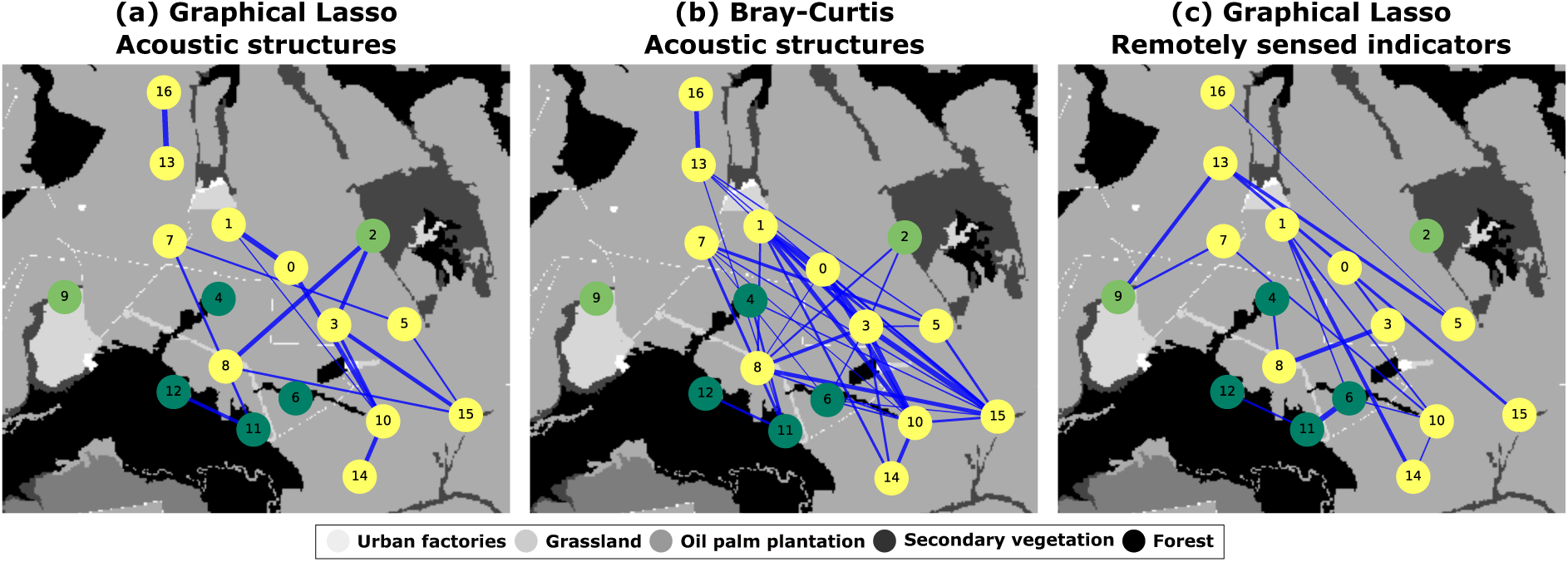
Comparative analysis of ecological heterogeneity identification models. Each node represents a distinct geographical site, with the color of the node indicating the type of land cover: yellow for oil palm plantations, light green for secondary vegetation, and dark green for forest. Edges represent the inferred similarities among sites. (a)Graphical Lasso model using acoustic structures (b) Bray-Curtis model using acoustic structures. (c) Graphical Lasso model with satellite indices. Coverage-type information was not utilized in computing the models.

#### 3.1.3. Bray-Curtis dissimilarity graph

Using the normalized sonotype occurrence matrix, we calculated the Bray-Curtis dissimilarity to estimate the similarities and relationships among geographical locations. The result is a 17 × 17 distance matrix with values ranging from 0 to 1, where values closer to 0 indicate higher similarity.

To visualize this distance matrix, we represented it as a graph. Each node in the graph corresponds to a sampled site, placed according to its geographic location. The color of each node represents the type of coverage for that site. The color significance is the same as in the Graphical Lasso network, and the links in the graph were determined based on the values in the dissimilarity matrix, where values below 0.4 were used for establishing connections.

The Bray-Curtis dissimilarity graph (see Figure 5 (b)) displays a generally homogeneous pattern. This can be attributed to the similarity-based connections between the various forest areas, secondary vegetation zones, and oil palm plantations. It shares some strong connections with the Graphical Lasso graph, particularly to nodes representing oil palm plantations ((0-1),(0-10),(10-14),(13-16)). Moreover, this result shows similar patterns to the Glasso graph representation, where forest sites 11–12 are connected but not connected with other forest places, sites 2–8 and 2-3 are also connected, and site 9 has no connections as in Glasso.

The Spearman correlation coefficient between the Graphical Lasso and the Bray-Curtis graphs is 0.55, indicating a moderately positive correlation. These two graphs share 12 common edges. The Graphical Lasso graph can be seen as a sub-graph of the Bray-Curtis graph, as it contains 78% of the links present in the Glasso model. However, its sparsity is the key advantage of the Graphical Lasso model over the Bray-Curtis dissimilarity approach. The Graphical Lasso tends to assign zero weights to dissimilar connections, resulting in a graph with more distinct patterns that are easier to analyze and are closer to what is expected by the experts.

#### 3.1.4. Remotely sensed indicators

On the other hand, we estimated a Graphical Lasso model using non-acoustic input features, specifically, a matrix of median values of coverage indices generated from satellite images (see Table 1). We obtained a 17 × 17 precision matrix describing similarities among the 17 locations. For this model, the optimal alpha parameter (corresponding to regularization *λ*) found through cross-validation was 0.18. To obtain a sparse graph representation, we retained precision matrix values above 0.7, following the same construction process described previously for the acoustic Graphical Lasso model.

It is possible to see in Figure 5c that the resulting graph contains several links among the geographical sites, suggesting a predominantly homogeneous landscape. The primary connections observed are between the oil palm plantations and the other two land cover types. Spearman correlation coefficient between acoustic Glasso and spectral indices Glasso is 0.11, indicating a non-correlation, evident in the Figure 5 where only 3 links are shared: (1-10), (10-14), and (11-12).

In this satellite data analysis, interconnected forest regions reveal analogous patterns within the imagery as in the case of nodes (11-12) (11-6), a trend also observed among the oil palm plantations. Here, the Graphical Lasso highlights the uniformity of their spectral signatures. The connections between sites, including (4-8), (6-10), and (1-6), uncover a pattern of uniformity in the satellite imagery that bridges the oil palm plantations with the small forest fragments. These linkages highlight that the captured features for these land covers are similar, exhibiting poor variation in the spectral data. Similarly, the links involving node 9 with additional oil palm locales imply a shared spectral characteristic. In contrast, node 2 is different, exhibiting unique patterns that diverge from other land covers, suggesting variations in vegetation values taken by the satellite. This starkly contrasts the acoustic Graphical Lasso model, where node 2 aligns with other nodes in acoustic patterns, while node 9 appears isolated.

Acoustic features provide an advantage when it is challenging to discern between land cover types and thus understand the heterogeneity among sites. It leverages the biophony patterns found at each location to identify similarities between sites, serving as a complementary tool to spectral indices analysis. Decomposing the landscape into acoustic entities allows for identifying the biophony patterns of each site, revealing similarities between the patterns across different locations.

#### 3.1.5. Acoustic Heterogeneity analysis and link interpretation

Building on the Graph Lasso model’s identification of acoustic similarities among different geographical sites, we aimed to understand the nature of these connections by analyzing the shared acoustic structures and sonotypes and their frequency-time information. This visualization depicts the frequency and timing of the most representative sonotypes for three site pairs: a pair whose connection is thicker (2–8), a connected pair (11–12), and a pair with no connection (4–9). Each dot, color-coded for individual sonotypes, is plotted against the time of detection and its peak frequency in addition to each acoustic time pattern.

Figure 6 presents a ‘soundscape visualization’ where we can see the distribution of common sonotypes, their frequency information, and acoustic time patterns across pairs of sites. Figures 6a and b display sites connected by the acoustic Graph Lasso model. For instance, sites 2-8, representing different types of cover, exhibit similar acoustic patterns related to shared sonotypes in the soundscape. A similar trend in sonotype activity in the daytime is observable in the temporal acoustic pattern, with peaks during sunrise and dusk within the 2–7 kHz range. These peaks are potentially linked to animal calls from those frequency bands, such as bird species, during those times.

**Figure 6.**
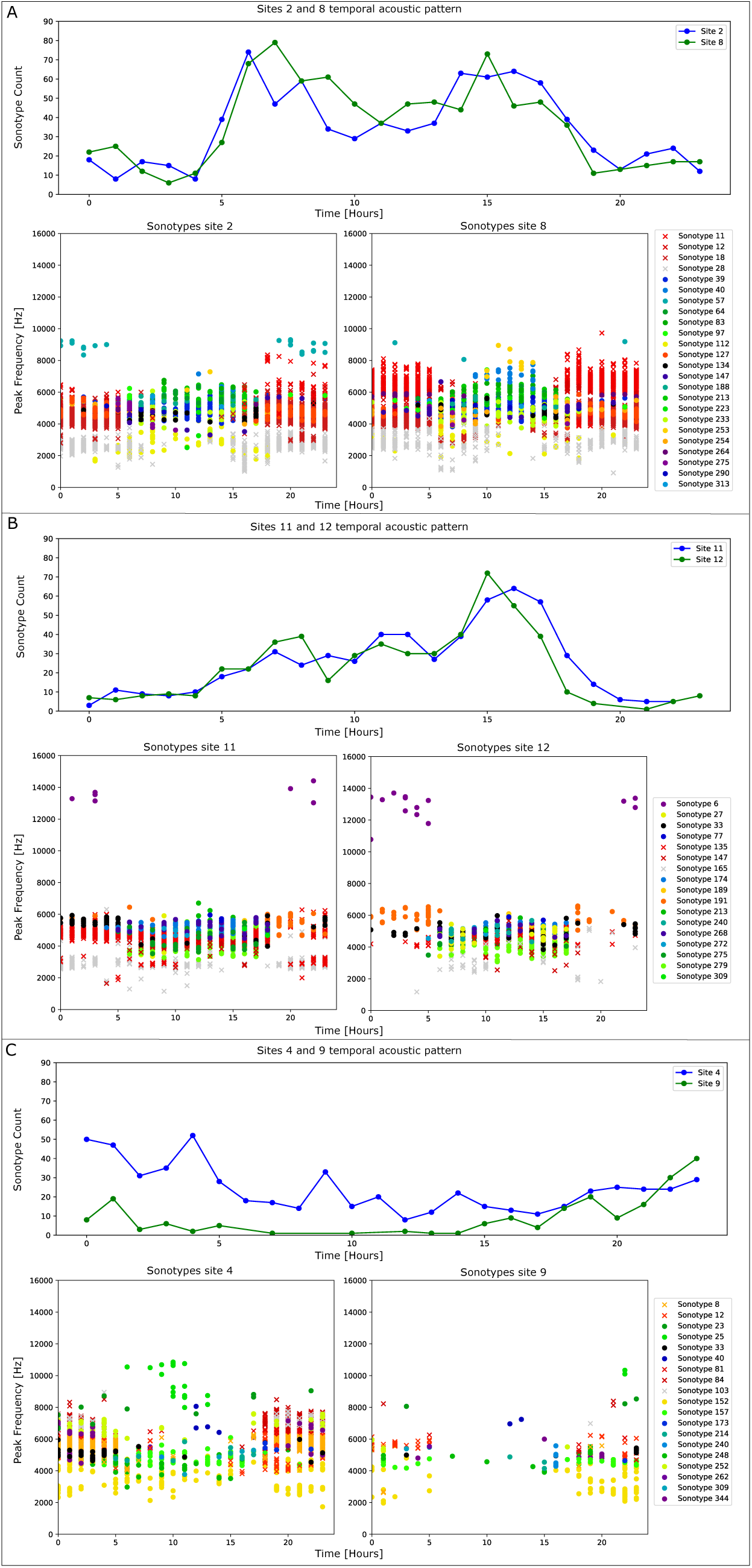
Visualization of acoustic patterns of representative sonotypes in three different site pairs. The upper chart plot presents the temporal activity of sonotypes, with sonotype activity on the vertical axis and time of day on the horizontal axis. The lower scatter plots map sonotype frequencies, with each sonotype represented by either a dot or a cross, each color-coded according to the specific sonotype indicated in the legend. The vertical axis represents peak frequency (Hz), and the horizontal axis represents time (hours). This visualization captures the diversity and temporal distribution of sonotypes within the soundscape of each site pair.

Connections between sites 11 and 12, depicted in Figure 6b and associated with forested areas, exhibit a similar acoustic pattern as observed in case a. There is a pronounced peak in activity at dusk, with frequencies ranging from 3-6 kHz. Additionally, high-frequency detections (above 12 kHz) found at both locations suggest a comparable nocturnal pattern, possibly related to insect stridulation or low-frequency bat calls.

Figure 6c illustrates the acoustic signatures of site pairs that Glasso did not connect, suggesting these are acoustically heterogeneous. These pairs of sites present particularities, such as the fact that they were isolated. It is possible to identify the difference in the distribution of the sonotypes and the acoustic time pattern, even showing the similar ones between both sites. In both cases, a low number of occurrences were present.

Interestingly, in the case of node 4, despite representing the same type of land cover as nodes 11-12, there is no acoustic connection between them or with other forests (node 6). This indicates that, even though they share the same coverage type, a barrier, such as oil palm cultivation, produces remarkable differentiation in their acoustic structures. Therefore, they possibly do not share sonotypes or differ significantly in occurrences, which leads to the assumption that there are animals that cannot move from one forest to another due to this physical barrier.

This type of representation, where the sonotype frequency bands are discernible, opens up the possibility of linking specific sonotypes to particular species, offering a valuable direction for further ecological investigation.

### 3.2. Case 2: Puerto Wilches temporal analysis

By utilizing the results obtained from the acoustic tool for automatic sonotype identification in the Puerto Wilches dataset, we estimated the acoustic structures for each time frame, dawn (05-8), day (08-16), dusk (16-20), and night (20-05), within each site to generate graphs that illustrate the intersite connections. This process yielded four matrices with dimensions of 17 × 292. We used the normalized version of these matrices to generate the Graphical Lasso model for each time frame, following an identical procedure to the one presented in case study 1.

Figure 7 shows the four generated graphs, each representing a different time frame. These graphs illustrate the dynamic changes in acoustic patterns, resulting in variations in similarities and connections among the sites. Furthermore, they reveal significant patterns related to the same land cover types.

**Figure 7.**
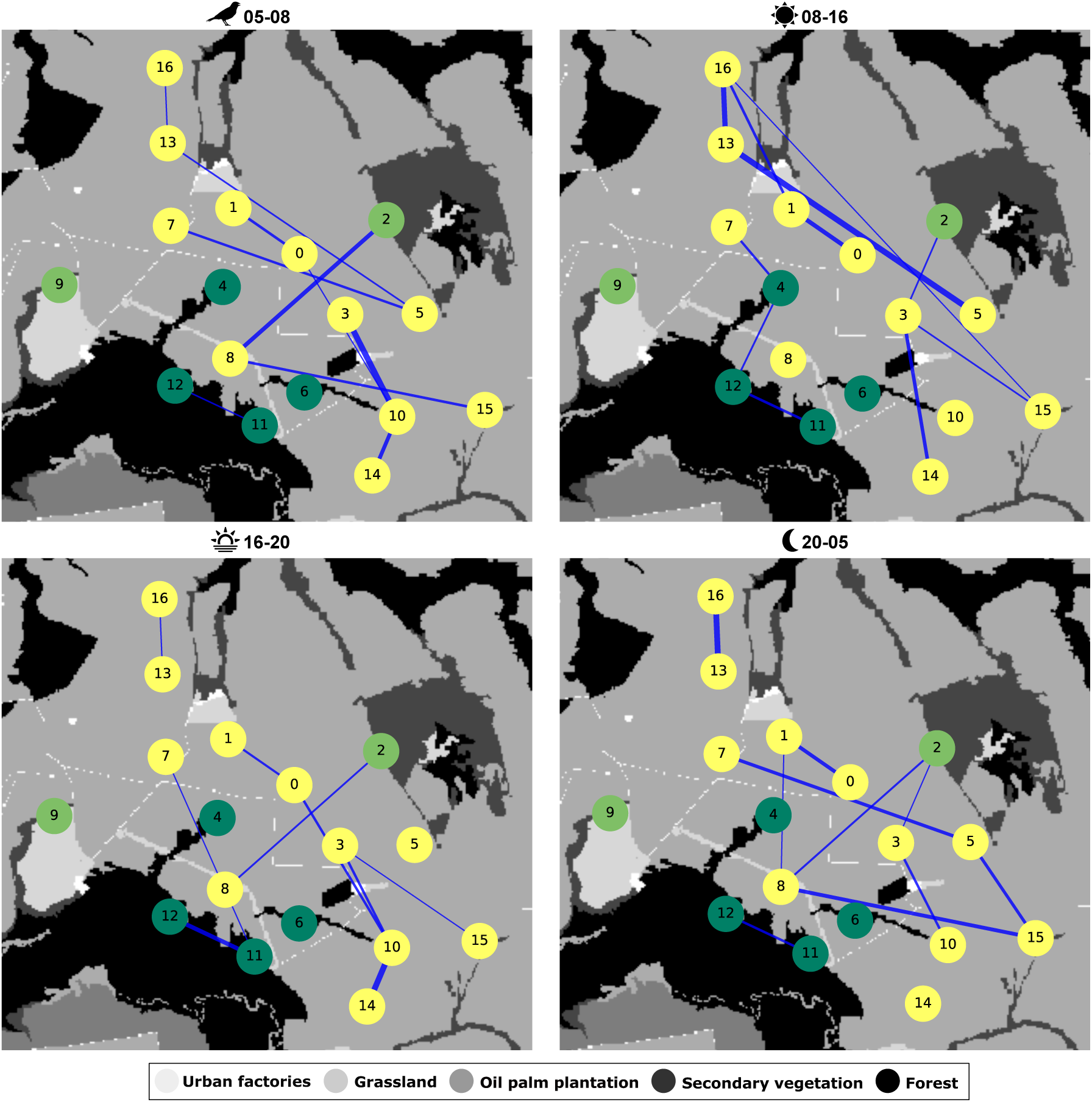
Graphical Lasso model for different time frames displaying similarities between sites using an edge. Nodes are located according to their geographical location, and their color represents their cover land type: yellow, oil palm plantation; light green, secondary vegetation; and dark green, forest. land cover-type information was not utilized in computing the models.

The graphical representation in Figure 6 sheds considerable light on these acoustic connections. Particularly for nodes 2 and 8, as depicted in Figure 6A, a pattern emerges where sonotypes are predominantly active during the early morning and dusk. However, nodes 4 and 9 are still not connected at different times of the day.

Figure 8 displays the network density across four distinct periods and the all-day graph generated in case study 2. Network density is a measure that indicates the proportion of actual connections present in the graph compared to the maximum possible connections. In this case, the graph shows variability in network density, with the day period exhibiting the highest density among the individual periods. This could indicate a surge in acoustic activity or events that facilitate or require increased communication or connectivity among the sites. Alternatively, this higher density may reflect a generally greater acoustic similarity during the daytime, potentially driven by a phenomenon of subtractive homogenization within acoustic communities active at this time. Given the direct relationship between sonotype composition and acoustic community structure and considering the historical landscape transformations in the region—marked by extensive deforestation and habitat fragmentation—this homogenization could result from a decline in the diversity of sound-emitting species, leading to the higher compositional similarity among sites. Furthermore, given our study scale, homogeneity in acoustic communities could be maintained as species active during the daytime could easily move or communicate between close sites, thus increasing the acoustic similarity.

**Figure 8.**
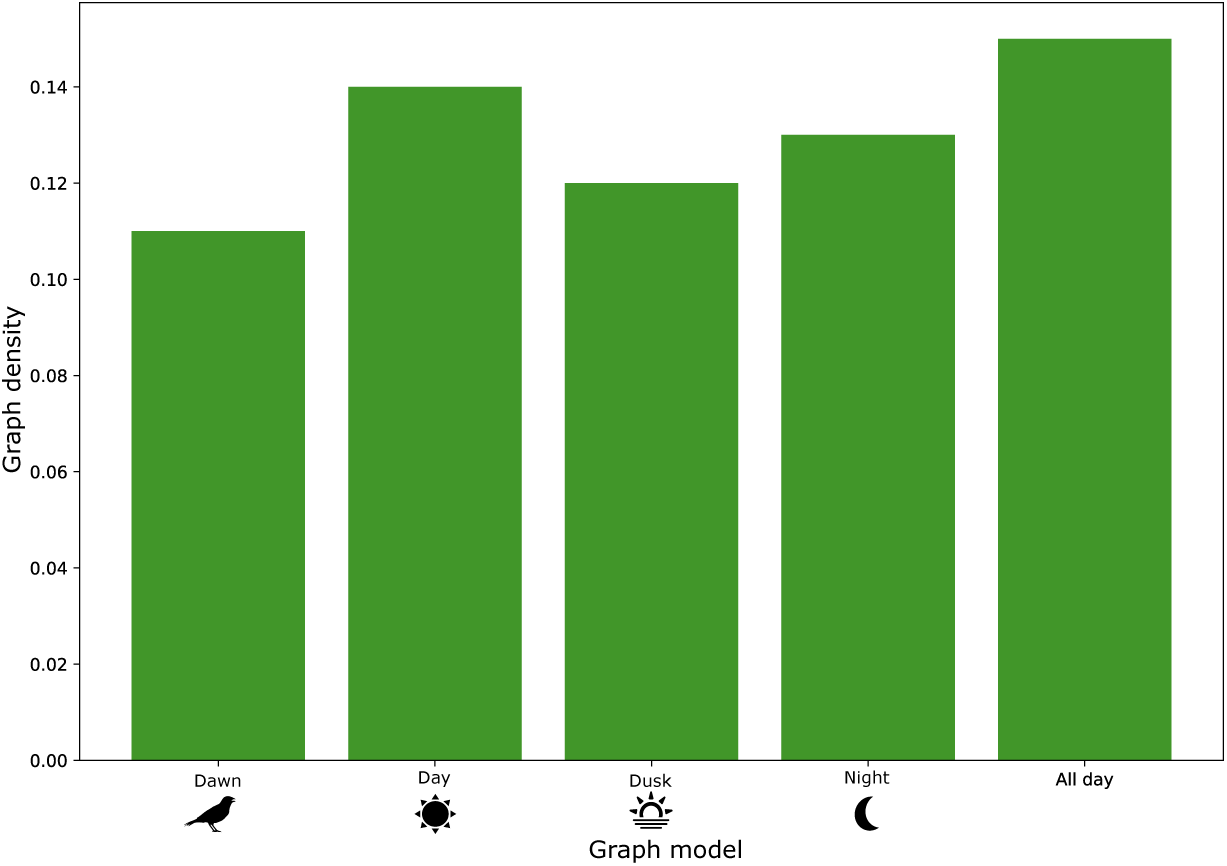
Network density across different periods and the all-day graph. This chart illustrates the network density for four separate time periods and an all-day graph, presented in case study 2. Each bar represents the density value, indicating the proportion of actual connections relative to the total possible connections within the network.

The density in the all-day graph is notably higher than in any individual period, highlighting the cumulative nature of connections throughout the day. This elevated density demonstrates that the all-day graph combines the patterns shown in the temporal graphs, reflecting an overall network structure that is highly interconnected when all connections are considered together. This suggests an ecologically integrated and dynamically adaptive network where different nodes may play crucial roles at different times, contributing to a diverse and interconnected landscape throughout the day. Moreover, the temporal graphs allow for identifying specific patterns and interactions, indicating that individual period analyses are crucial for understanding nuanced ecological and soundscape dynamics.

## 4. Discussion

In this study, we presented a novel unsupervised framework to assess landscape heterogeneity using passive acoustic monitoring with network inference analysis. Our approach does not require prior labels for animal calls or coverage types, making it versatile for various ecological applications. This method enables the differentiation of geographical sites and landscapes based on acoustic structures composed of sonotypes. These sonotypes are distinctive sound patterns that occupy specific frequency and time intervals and can be linked to species calls (Guerrero et al., 2023). Utilizing the network inference data and sonotype-derived time-frequency information, we performed two graphical representations to elucidate connections among site pairs, revealing the spectral-temporal characteristics driving these similarities.

Our findings highlight the performance of acoustic structures as input features for the Graphical Lasso (Glasso) model over traditional methods, such as Bray-Curtis dissimilarity, in identifying similarity patterns. The Glasso model’s sparse architecture succeed at revealing the intricate relationships within complex acoustic landscapes. Moreover, because these acoustic structures are composed of sonotypes, they offer details of how the soundscape is conformed, thereby providing a more nuanced analysis. This advantage becomes particularly evident when compared to satellite indices, which struggle with accurately distinguishing areas in transitional states or small geographic extents despite their widespread use for land cover identification. In these cases, our acoustic analysis presents a viable complement, offering insights with a different resolution than what spectral indices can achieve, potentiating the results for identifying landscape heterogeneity.

Bormpoudakis et al. (2013) demonstrated that ambient sound is spatially heterogeneous and directly associated with habitat structure, indicating habitat-specific acoustic signatures and each habitat type has a unique soundscape. This underscores the importance of using acoustic characteristics to understand and identify similarities between what is a habitat for sonoriferous species. Our study extends this idea by decomposing the soundscape into sonotypes, which allows for a detailed analysis of acoustic habitats and their heterogeneity. The strategic decomposition of the soundscape into sonotypes illustrates the method’s advantage in understanding ecological interactions, not just through similarity analysis but also by understanding the underlying patterns and information driving these similarities. This approach marks a significant departure from conventional methods reliant solely on correlation, distance analyses, and dendrograms, offering a more nuanced interpretation of ecological dynamics and the possibility of associating the identified acoustic patterns with species calls.

Analyzing the Puerto Wilches dataset across distinct daily periods revealed temporal dependencies, validating the method’s responsiveness to biological rhythms, consistent with previous studies (Deichmann et al., 2017; Sánchez-Giraldo et al., 2021; Rendon et al., 2022; Barbaro et al., 2022). Acoustic profiles in oil palm plantations showed different variations in graph structure across time frames, potentially reflecting species dynamics and anthropogenic influences shaped by structured agricultural practices. Such insights into the temporal dynamics of sonotypes within these human-modified landscapes are critical for developing targeted conservation strategies that consider the biological and anthropogenic factors shaping these ecosystems. Future research should consider the analysis of the landscape at different stages of the day. This could support the interpretation of models, as in the case of identifying species in (Jeantet and Dufourq, 2023).

Despite these strengths, the method has certain limitations. The Glasso model requires threshold selection, potentially hiding the direct interpretation of significant acoustic features, a common issue with other graph inference methods (Brugere et al., 2018). Consequently, future research should focus on developing inference models that minimize or eliminate subjective thresholding, enhancing intuitive ecological interpretation without auxiliary analyses such as distance computations.

Moreover, accurate ecological interpretation from acoustic data strongly depends on dataset quality and preprocessing to ensure that analyzed acoustic signals predominantly represent biophony. Our methodology can potentially capture non-biophonic components depending on dataset conditions. Thus, careful initial dataset curation is crucial. In our dataset, anthropogenic disturbances were minimal and unlikely to influence our conclusions significantly. Nevertheless, scenarios with higher human activity could generate sonotypes, and they will still be informative, as they would contribute meaningfully to the similarity patterns and network structure, capturing a broader view of the acoustic landscape. However, cautious interpretation may be necessary to ensure accurate ecological insights.

It is also important to highlight that our sonotype-based approach does not directly imply taxonomic identification or ecological similarity traditionally obtained through species lists or taxonomic measures. Instead, this method offers a complementary, unsupervised, cost-effective approach to assessing biological similarity using acoustic data. This analytical advantage not only provides an initial indication of acoustic similarity but also suggests biological similarity among sites. Based on the positive link between acoustic similarity and landscape connectivity proxies (e.g., (Burivalova et al., 2019; Hayashi et al., 2020)), our approach could also serve as a potential complementary tool for early assessment and monitoring of connectivity or fragmentation measures at diverse spatial scales, aiding management, and conservation decisions, especially when traditional ecological surveys are logistically or financially challenging.

This work not only highlights the potential of acoustic monitoring in ecological network inference but also points towards the need for more interpretable and direct analysis methods. Doing so will pave the way for future studies to further refine and expand upon acoustic data in ecological research, enhancing our understanding of biodiversity and ecosystem dynamics.

## Acknowledgments

This work was supported by Universidad de Antioquia - CODI and Alexander von Humboldt Institute for Research on Biological Resources. [code project:2020-33250].Part of this material is based upon work supported by the National Science Foundation under Grants #2213568, and Rice University Creative Ventures: Sustainable Futures Fund. Data from Puerto Wilches were funded by Universidad de Antioquia, SGI, and Ecopetrol under contract FOGR09.

## Authors’ Contributions

**Maria J. Guerrero** conceptualization, methodology, algorithms, writing the manuscript; **Camilo Sánchez** and **Víctor M. Martínez-Arias** data acquisition, model validation, model interpretation, and ecology conceptualization; **Cé sar A. Uribe** graph inference conceptualization, methodology, supervised the research; **Claudia Isaza** conceptualization, supervision, methodology, validation, and project administration. All authors contributed to the drafts and gave the final approval for publication.

## Data Availability

The data and codes used for all case studies, comparison methods, and evaluation cases are available on this link: Data and codes here.

## Appendix A

### Evaluation case: soundscape simulation

Artificial acoustic structures were created based on the expected acoustic behavior patterns at sites, mirroring a practice in graph inference where a proposed approach is illustrated using synthetic datasets. These simulated acoustic structures aim to reflect different scenarios in a natural environment (e.g., the same composition of sonotypes, similar occurrences, or different composition of sonotypes and different occurrences), enabling detailed, case-by-case analysis with a clear understanding of expected outcomes.

An 11 × 10 matrix was created to represent 11 artificial sites, each associated with 10 sonotypes and their corresponding number of occurrences. Figure A.1 presents the generated acoustic structures, where each color represents an artificial coverage type, and all radial plots show the 10 sonotypes and their occurrences in each simulated site.

**Figure A.1.**
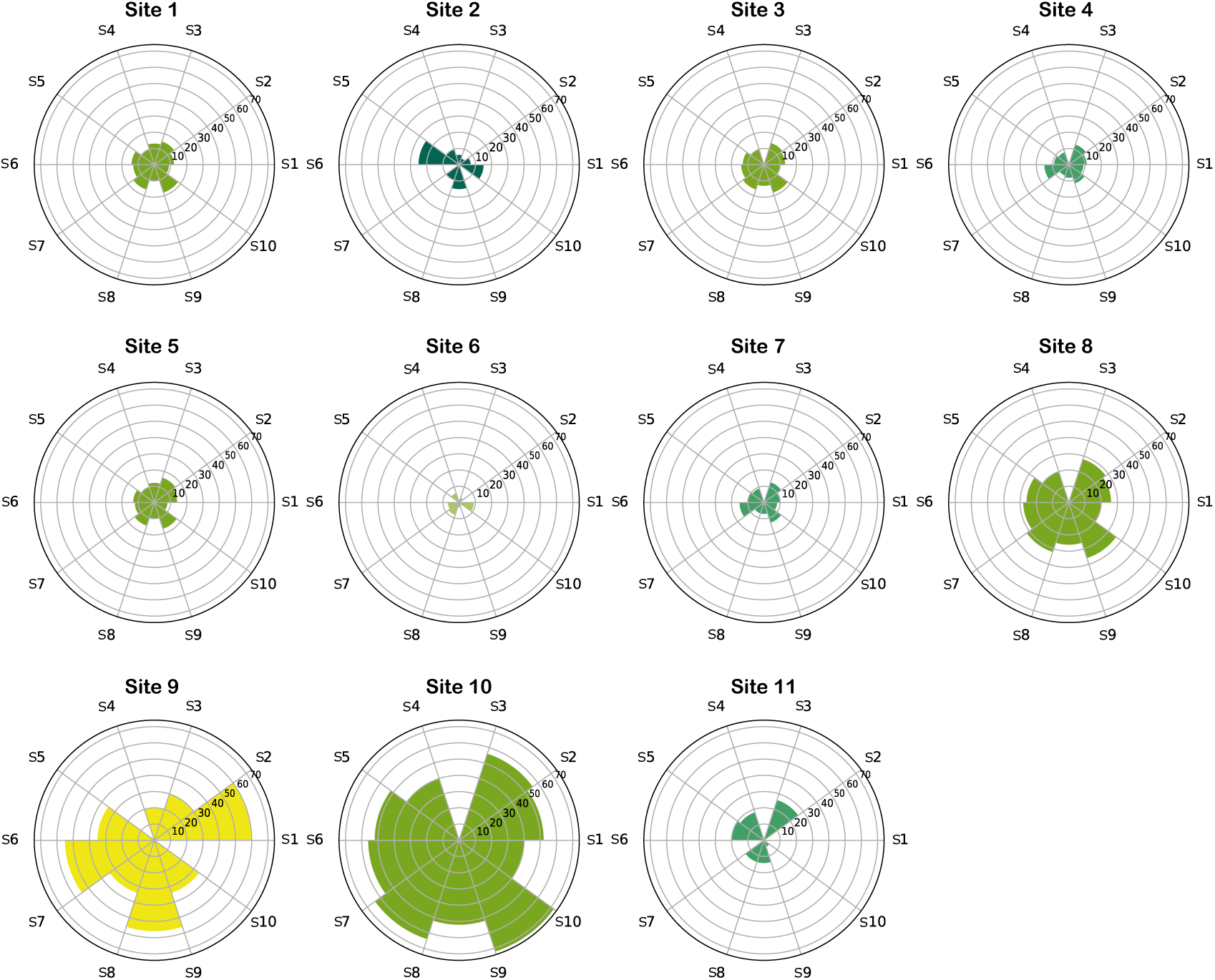
Simulated acoustic structures. Each structure is composed of the number of occurrences of its sonotypes. Structures with the same color (e.g., 4, 7, 11) represent similar land cover types (artificially).

Figure A.2 depicts the expected graph based on the simulated acoustic structures, where the color of each node (site) indicates whether it represents an artificial coverage type, the same if nodes (sites) have the same color, and different otherwise. Based on the similarity of the structures, we would expect that sites with the same or similar structures are connected. The width of the edges in the graph represents the strength of the connection between sites, given by the value of the precision matrix. Subsequently, we utilize the corresponding adjacency matrix to calculate the confusion matrix, which evaluates the performance of the graph models generated with different methods.

The performance of this evaluation case was assessed using the accuracy metric, utilizing a confusion matrix to contrast the links inferred by the evaluated methods against the ground truth. This involves the comparison between the adjacency matrix **A** ∈ R*^m^*^×*m*^ of the expected graph and the adjacency matrix **A**^′^ ∈ R*^m^*^×*m*^ of the resultant graph. To calculate accuracy, we accurately identify true positives (TP), false positives (FP), true negatives (TN), and false negatives (FN) using these matrices. Accuracy is the proportion of correctly identified links (both true positives and true negatives) relative to the total number of evaluated links. The formula for accuracy is given by:

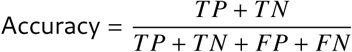

Where:

- *TP* (*True Positives*) are the links present in both **A** and **A**^′^.
- *TN* (*True Negatives*) are the links absent in both **A** and **A**^′^.
- *FP* (*False Positives*) are the links not present in **A** but present in **A**^′^.
- *FN* (*False Negatives*) are the links present in **A** but not in **A**^′^.

These values are calculated by comparing each corresponding element of matrices **A** and **A**^′^. An element *a_i j_* in matrix **A** and the corresponding element 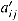 in **A**^′^ contribute to *TP* if both are 1, to *TN* if both are 0, to *FP* if *a_i j_* = 0 and 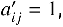 and to *FN* if *a_i j_* = 1 and 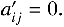

The artificial structures created (matrix of occurrences of artificial sonotypes) for this case are shared at: (Available on Figshare when the manuscript is accepted)

### Appendix A.1. Results

The simulated acoustic structures were analyzed using the Graphical Lasso model and Bray-Curtis dissimilarity to estimate the homogeneity/heterogeneity among sites and compare both techniques first. Considering the ground truth (Figure A.2a), these results were evaluated by calculating the accuracy based on the confusion matrix for both methods.

#### Appendix A.1.1. Graphical Lasso model

Graph Lasso method processes the normalized matrix of acoustic structures of each location, in this case, the 11 × 10 artificial matrix (11 sites and 10 sonotypes), as input values. This method employs an automated cross-validation process to determine the optimal alpha value, which was found to be 0.02.

As a result, an 11×11 precision matrix was obtained; this matrix reflects the relationships among the sites under study. Then, the values of the precision matrix are considered in the construction of the resulting graph, where a threshold value (*th* = 0.5) was necessary to establish the relevant connections.

Figure A.2b presents the resulting graph based on the precision matrix generated by Graphical Lasso. Here, each color node (four types of green and yellow) represents a simulated type of coverage (5 different types), and edge width represents a higher value in the precision matrix that indicates a stronger connection between sites. The accuracy of the estimation of this graph was 98% (see confusion matrix in table A.1), where only one edge was missing (7-11) to have a perfect estimation according to what the ground truth presents.

#### Appendix A.1.2. Bray-Curtis dissimilarity analysis

To generate a graph based on the Bray-Curtis dissimilarity, it was necessary to use the simulated acoustic structure matrix. As well as Graphical Lasso, this matrix was normalized. Then, an 11 × 11 Bray-Curtis matrix was estimated. This matrix has values between 0 and 1 that indicate how similar or different the sites are, where values close to 0 indicate similarity. Using this dissimilarity matrix, we obtain a graph representation that connects all the more similar places according to their small values. Figure A.2c shows a graph construction based on the Bray-Curtis dissimilarity. In this case, the accuracy was 78% (see confusion matrix in table A.1) concerning the ground truth network. However, connections represented as strong in the ground truth graph, such as (1-5) and (7-4), and isolated nodes, such as 6 and 11, were presented here. It is important to highlight that in this case, it was possible to determine the performance of both comparison models due to the ground truth (see Figure A.2a) obtained from the simulated acoustic structures (see Figure A.1).

**Figure A.2.**
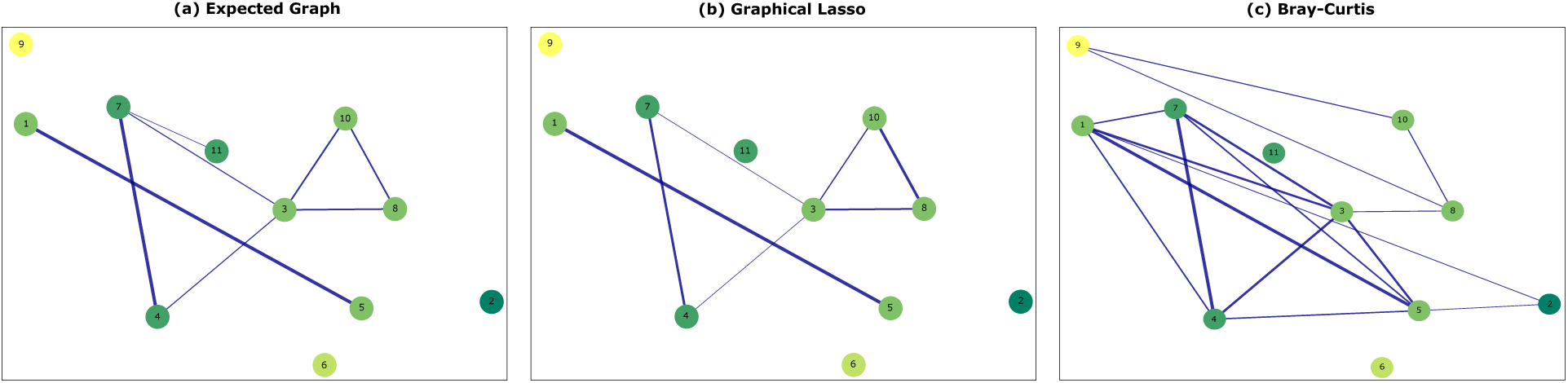
Graph analysis of simulated acoustic structures across 11 different sites. Each node color represents a land cover type, while the width of the links indicates the strength of the connection between the locations. Artificial sites with similar acoustic structures are expected to have strong connections. (a) The expected graph is derived from the simulated acoustic structures. (b) The resultant graph from Graph Lasso was applied to the simulated acoustic structures. (c) Bray-Curtis dissimilarity-based graph generated from the simulated acoustic structures.

**Table A.1:**
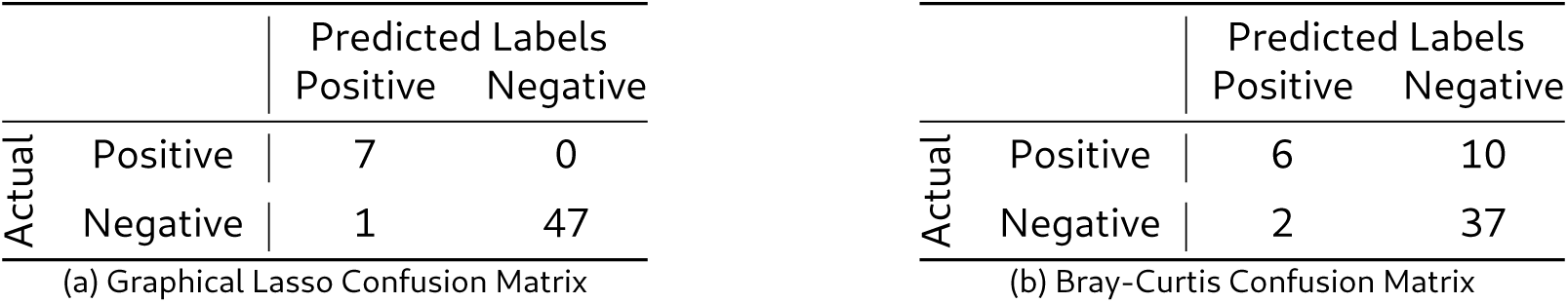
Confusion matrices for Graphical Lasso and Bray-Curtis results concerning the ground truth.

## Footnotes

1 http://www.humanconnectomeproject.org/

## References

Acevedo, M. and Villanueva-Rivera, L. (2006). Using automated digital recording systems as effective tools for the monitoring of birds and amphibians. Wildlife Society Bulletin 34: 211–214, doi:10.2193/0091-7648(2006)34[211:UADRSA]2.0.CO;2.

Al-Wassai, F. A. and Kalyankar, N. (2013). Major limitations of satellite images. *arXiv preprint arXiv*:*1307.2434*.

Alcocer, I., Lima, H., Sugai, L. and Llusia, D. (2022). Acoustic indices as proxies for biodiversity: a meta-analysis. Biological Reviews 97, doi:10.1111/brv.12890.

Ali, A. M., Darvishzadeh, R., Skidmore, A. K. and van Duren, I. (2017). Specific leaf area estimation from leaf and canopy reflectance through optimization and validation of vegetation indices. Agricultural and Forest Meteorology 236: 162–174, 10.1016/j.agrformet.2017.01.015.

Apoux, F., Miller-Viacava, N., Ferrière, R., Dai, H., Krause, B., Sueur, J. and Lorenzi, C. (2023). Auditory discrimination of natural soundscapes. The Journal of the Acoustical Society of America 153: 2706, doi:10.1121/10.0017972.

Barbaro, L., Sourdril, A., Froidevaux, J. S., Cauchoix, M., Calatayud, F., Deconchat, M. and Gasc, A. (2022). Linking acoustic diversity to compositional and configurational heterogeneity in mosaic landscapes. Landscape Ecology doi:10.1007/s10980-021-01391-8.

Bedoya, C., Isaza, C., Daza, J. M. and López, J. D. (2014). Automatic recognition of anuran species based on syllable identification. Ecological Informatics 24: 200–209, doi:10.1016/j.ecoinf.2014.08.009.

Bedoya, C., Isaza, C., Daza, J. M. and López, J. D. (2017). Automatic identification of rainfall in acoustic recordings. Ecological Indicators 75: 95–100, doi:10.1016/j.ecolind.2016.12.018.

Bellisario, K., Broadhead, T., Savage, D., Zhao, Z., Omrani, H., Zhang, S., Springer, J. and Pijanowski, B. (2019). Contributions of mir to soundscape ecology. part 3: Tagging and classifying audio features using a multi-labeling k-nearest neighbor approach. Ecological Informatics 51, doi:10.1016/j.ecoinf. 2019.02.010.

Berra, E. and Gaulton, R. (2021). Remote sensing of temperate and boreal forest phenology: A review of progress, challenges and opportunities in the intercomparison of in-situ and satellite phenological metrics. Forest Ecology and Management 480: 17, doi:10.1016/j.foreco.2020.118663.

Bhushan, N., Mohnert, F., Sloot, D., Jans, L., Albers, C. and Steg, L. (2019). Using a gaussian graphical model to explore relationships between items and variables in environmental psychology research. Frontiers in Psychology 10: 1050, doi:10.3389/fpsyg.2019.01050.

Bicudo, T., Llusia, D., Anciães, M. and Gil, D. (2023). Poor performance of acoustic indices as proxies for bird diversity in a fragmented amazonian landscape. Ecological Informatics 77: 102241, doi: 10.1016/j.ecoinf.2023.102241.

Bormpoudakis, D., Sueur, J. and Pantis, J. (2013). Spatial heterogeneity of ambient sound at the habitat type level: Ecological implications and applications. Landscape Ecology 28: 495–506, doi:10.1007/s10980-013-9849-1.

Bray, J. R. and Curtis, J. T. (1957). An ordination of the upland forest communities of southern wisconsin. Ecological Monographs 27: 325–349, doi:10.2307/1942268.

Brugere, I., Gallagher, B. and Berger-Wolf, T. (2018). Network structure inference, a survey: Motivations, methods, and applications. ACM Computing Surveys 51, doi:10.1145/3154524.

Burivalova, Z., Purnomo, P., Wahyudi, B., Boucher, T., Ellis, P., Truskinger, A., Towsey, M., Roe, P., Marthinus, D., Griscom, B. and Game, E. (2019). Using soundscapes to investigate homogenization of tropical forest diversity in selectively logged forests. Journal of Applied Ecology 56, doi:10.1111/1365-2664.13481.

Castro-Ospina, A., Solarte-Sanchez, M., Vega-Escobar, L., Isaza, C. and Martinez-Vargas, J. (2024). Graph-based audio classification using pre-trained models and graph neural networks. Sensors 24: 2106, doi:10.3390/s24072106.

Castro-Ospina, A. E., Rodriguez-Buritica, S., Rendon, D., Velandia-García, M., Isaza, C. and Martínez-Vargas, J. (2023). Identification of Tropical Dry Forest Transformation from Soundscapes Using Supervised Learning. 173–184, doi:10.1007/978-3-031-32213-613.

Correa Ayram, C., Díaz, J., Etter, A., Ramírez, W. and Corzo, G. (2019). El cambio en la huella espacial humana como herramienta para la toma de decisiones en la gestión del territorio.

Deichmann, J., Hernandez-Serna, A., Delgado Cornejo, J. A., Campos Cerqueira, M. and Aide, T. M. (2017). Soundscape analysis and acoustic monitoring document impacts of natural gas exploration on biodiversity in a tropical forest. Ecological Indicators 74, doi:10.1016/j.ecolind.2016.11.002.

Dufourq, E., Durbach, I., Hansford, J., Hoepfner, A., Ma, H., Bryant, J., Stender, C., Li, W., Liu, Z., Chen, Q., Zhou, Z. and Turvey, S. (2020). Automated detection of hainan gibbon calls for passive acoustic monitoring. Remote Sensing in Ecology and Conservation doi:10.1101/2020.09.07.285502.

Dumyahn, S. L. and Pijanowski, B. C. (2011). Soundscape conservation. Landscape Ecol 26: 1327–1344, doi:10.1007/s10980-011-9635-x.

Equator Studios (Accessed: 2023). How to Calculate Slope in QGIS. https://equatorstudios.com/how-to-calculate-slope-in-qgis.

Farina, A. (2022). Principles and Methods in Landscape Ecology: An Agenda for the Second Millennium. Switzerland: Springer Nature Switzerland AG, doi:10.1007/978-3-030-96611-9, in Principles and Methods in Landscape Ecology.

Fitzgerald, G., Rodriguez, D. and O’Leary, G. (2010). Measuring and predicting canopy nitrogen nutrition in wheat using a spectral index—the canopy chlorophyll content index (ccci). Field Crops Research 116: 318–324, 10.1016/j.fcr.2010.01.010.

Friedman, J., Hastie, T. and Tibshirani, R. (2008). Sparse inverse covariance estimation with the graphical lasso. Biostatistics (Oxford, England) 9: 432–41, doi:10.1093/biostatistics/kxm045.

Fu, X., Seo, E., Clarke, J. and Hutchinson, R. A. (2021). Link prediction under imperfect detection: Collaborative filtering for ecological networks. IEEE Transactions on Knowledge & Data Engineering 33: 3117–3128, doi:10.1109/TKDE.2019.2962031.

Fuller, S., Axel, A., Tucker, D. and Gage, S. (2015). Connecting soundscape to landscape: Which acoustic index best describes landscape configuration? Ecological Indicators 58: 207–215, doi: 10.1016/j.ecolind.2015.05.057.

Gao, B. C. (1996). Ndwi—a normalized difference water index for remote sensing of vegetation liquid water from space. Remote sensing of environment 58: 257–266.

Gao, J., Sharma, R., Qian, C., Glass, L., Spaeder, J., Romberg, J., Sun, J. and Xiao, C. (2021). Stan: spatio-temporal attention network for pandemic prediction using real-world evidence. Journal of the American Medical Informatics Association 28, doi:10.1093/jamia/ocaa322.

Gasc, A., Pavoine, S., Lellouch, L., Grandcolas, P. and Sueur, J. (2015). Acoustic indices for biodiversity assessments: Analyses of bias based on simulated bird assemblages and recommendations for field surveys. Biological Conservation 191: 306–312, doi:10.1016/j.biocon.2015.06.018.

Gibb, R., Browning, E., Glover-Kapfer, P. and Jones, K. E. (2018). Emerging opportunities and challenges for passive acoustics in ecological assessment and monitoring. Methods in Ecology and Evolution 2019: 169–185, doi:10.1111/2041-210X.13101.

Gitelson, A. A. and Merzlyak, M. N. (1997). Remote estimation of chlorophyll content in higher plant leaves. International Journal of Remote Sensing 18: 2691–2697, doi:10.1080/014311697217558.

Guerrero, M. J., Bedoya, C. L., López, J. D., Daza, J. M. and Isaza, C. (2023). Acoustic animal identification using unsupervised learning. Methods in Ecology and Evolution 14: 1500–1514, doi: 10.1111/2041-210X.14103.

Gómez, W., Isaza, C. and Daza, J. (2018). Identifying disturbed habitats: A new method from acoustic indices. Ecological Informatics 45, doi:10.1016/j.ecoinf.2018.03.001.

Hagberg, A. A., Schult, D. A. and Swart, P. J. (2008). Exploring network structure, dynamics, and function using NetworkX. In Varoquaux, G., Vaught, T. and Millman, J. (eds), Proceedings of the 7th Python in Science Conference (SciPy2008). Pasadena, CA USA, 11–15.

Hayashi, K., Erwinsyah, Lelyana, V. D. and Yamamura, K. (2020). Acoustic dissimilarities between an oil palm plantation and surrounding forests: Analysis of index time series for beta-diversity in south sumatra, indonesia. Ecological Indicators 112, doi:10.1016/j.ecolind.2020.106086.

Huang, Y.-J., Lu, T.-P. and Hsiao, C. K. (2020). Application of graphical lasso in estimating network structure in gene set. Annals of Translational Medicine doi:10.21037/atm-20-6490.

Huete, A., Didan, K., Miura, T., Rodriguez, E., Gao, X. and Ferreira, L. (2002). Overview of the radiometric and biophysical performance of the modis vegetation indices. Remote Sensing of Environment 83: 195–213, doi:10.1016/S0034-4257(02)00096-2.

Jaeger, J. A. (2000). Landscape division, splitting index, and effective mesh size: new measures of landscape fragmentation. Landscape Ecology 15: 115–130, doi:10.1023/A:1008129329289.

Jeantet, L. and Dufourq, E. (2023). Improving deep learning acoustic classifiers with contextual information for wildlife monitoring. Ecological Informatics 77: 102256, doi:10.1016/j.ecoinf.2023.102256.

Jinru, X. and Su, B. (2017). Significant remote sensing vegetation indices: A review of developments and applications. Journal of Sensors 2017: 1–17, doi:10.1155/2017/1353691.

Kallimanis, A. S., Mazaris, A. D., Tsakanikas, D., Dimopoulos, P., Pantis, J. D. and Sgardelis, S. P. (2012). Efficient biodiversity monitoring: Which taxonomic level to study? Ecological Indicators 15: 100– 104, 10.1016/j.ecolind.2011.09.024.

Kriegler, F. J., Malila, W. A., Nalepka, R. F. and Richardson, W. (1969). Preprocessing transformations and their effects on multispectral recognition. Remote sensing of environment 97.

Lambin, E. and Meyfroidt, P. (2011). Global land use change, economic globalization, and the looming land scarcity. Proceedings of the National Academy of Sciences of the United States of America 108: 3465–72, doi:10.1073/pnas.1100480108.

LeBien, J., Zhong, M., Campos-Cerqueira, M., Velev, J. P., Dodhia, R., Ferres, J. L. and Aide, T. M. (2020). A pipeline for identification of bird and frog species in tropical soundscape recordings using a convolutional neural network. Ecological Informatics 59: 101113, doi:10.1016/j.ecoinf.2020.101113.

Legendre, P. and Legendre, L. (1998). Numerical Ecology. 1–853.

Li, R., Yuan, X., Radfar, M., Marendy, P., Ni, W., O’Brien, T. and Casillas-Espinosa, P. (2021). Graph signal processing, graph neural network and graph learning on biological data: A systematic review. IEEE Reviews in Biomedical Engineering PP: 1–1, doi:10.1109/RBME.2021.3122522.

Llusia, D. (2024). The limits of acoustic indices. Nature Ecology & Evolution 8, doi:10.1038/s41559-024-02348-1.

Mazumder, R. and Hastie, T. (2011). The graphical lasso: New insights and alternatives. Electronic Journal of Statistics 6, doi:10.1214/12-EJS740.

Montero, D., Aybar, C., Mahecha, M., Martinuzzi, F., Söchting, M. and Wieneke, S. (2023). A standardized catalogue of spectral indices to advance the use of remote sensing in earth system research. Scientific Data 10: 197, doi:10.1038/s41597-023-02096-0.

Ndao, B., Leroux, L., Gaetano, R., Diouf, A. A., Soti, V., Bégué, A., Mbow, C. and Sambou, B. (2021). Landscape heterogeneity analysis using geospatial techniques and a priori knowledge in sahelian agro-forestry systems of senegal. Ecological Indicators 125: 107481, 10.1016/j.ecolind.2021.107481.

Newbold, T., Hudson, L., Hill, S., Contu, S., Lysenko, I., Senior, R., Börger, L., Bennett, D., Choimes, A., Collen, B., Day, J., De Palma, A., Diaz, S., Echeverria-Londono, S., Edgar, M., Feldman, A., Garon, M., Harrison, M., Alhusseini, T. and Purvis, A. (2015). Global effects of land use on local terrestrial biodiversity. Nature 520: 45–50, doi:10.1038/nature14324.

Nolasco, I., Singh, S., Morfi, V., Lostanlen, V., Strandburg-Peshkin, A., Vidaña-Vila, E., Gill, L., Pamuła, H., Whitehead, H., Kiskin, I., Jensen, F., Morford, J., Emmerson, M., Versace, E., Grout, E., Liu, H., Ghani, B. and Stowell, D. (2023). Learning to detect an animal sound from five examples. Ecological Informatics 77: 102258, doi:10.1016/j.ecoinf.2023.102258.

Pedregosa, F., Varoquaux, G., Gramfort, A., Michel, V., Thirion, B., Grisel, O., Blondel, M., Prettenhofer, P., Weiss, R., Dubourg, V., Vanderplas, J., Passos, A., Cournapeau, D., Brucher, M., Perrot, M. and Duchesnay, E. (2011). Scikit-learn: Machine learning in Python. Journal of Machine Learning Research 12: 2825–2830.

Pijanowski, B., Farina, A., Gage, S., Dumyahn, S. and Krause, B. (2011). What is soundscape ecology? an introduction and overview of an emerging new science. Landscape Ecol 26: 1213–1232, doi: 10.1007/s10980-011-9600-8.

Potapov, P., Li, X., Hernandez-Serna, A., Tyukavina, A., Hansen, M. C., Kommareddy, A., Pickens, A., Turubanova, S., Tang, H., Silva, C. E., Armston, J., Dubayah, R., Blair, J. B. and Hofton, M. (2021). Mapping global forest canopy height through integration of gedi and landsat data. Remote Sensing of Environment 253: 112165, 10.1016/j.rse.2020.112165.

Powers, R. and Jetz, W. (2019). Global habitat loss and extinction risk of terrestrial vertebrates under future land-use-change scenarios. Nature Climate Change 9: 1, doi:10.1038/s41558-019-0406-z.

Radocaj, D., Obhodas, J., Jurišić, M. and Gasparovic, M. (2020). Global open data remote sensing satellite missions for land monitoring and conservation: A review. Land 9: 402, doi:10.3390/land9110402.

Ranciati, S., Roverato, A. and Luati, A. (2021). Fused Graphical Lasso for Brain Networks with Symmetries. Journal of the Royal Statistical Society Series C: Applied Statistics 70: 1299–1322, doi: 10.1111/rssc.12514.

Rendon, D., Rodriguez-Buritica, S., Sánchez-Giraldo, C., Daza, J. and Isaza, C. (2022). Automatic acoustic heterogeneity identification in transformed landscapes from colombian tropical dry forests. Ecological Indicators 140: 109017, doi:10.1016/j.ecolind.2022.109017.

Sousa-Lima, R., Fernandes, D., Norris, T. and Oswald, J. (2013). A review and inventory of fixed autonomous recorders for passive acoustic monitoring of marine mammals. Aquatic Mammals 39: 1–9, doi:10.1109/RIOAcoustics.2013.6683984.

Stowell, D. and Sueur, J. (2020). Ecoacoustics: acoustic sensing for biodiversity monitoring at scale. Remote Sensing in Ecology and Conservation 6: 217–219, doi:10.1002/rse2.174.

Sueur, J. and Farina, A. (2015). Ecoacoustics: the ecological investigation and interpretation of environmental sound. Biosemiotics 8: 493–502, doi:10.1007/s12304-015-9248-x.

Sugai, L. S. M., Silva, T. S. F., Ribeiro, J. W. and Llusia, D. (2019). Terrestrial passive acoustic monitoring: Review and perspectives. BioScience 69: 5–11, doi:10.1093/biosci/biy147.

Sánchez-Giraldo, C., Correa Ayram, C. and Daza, J. (2021). Environmental sound as a mirror of landscape ecological integrity in monitoring programs. Perspectives in Ecology and Conservation 19, doi:10.1016/j.pecon.2021.04.003.

Tscharntke, T., Tylianakis, J., Rand, T., Didham, R., Fahrig, L., Batary, P., Bengtsson, J., Clough, Y., Crist, T., Dormann, C., Ewers, R., Fründ, J., Holt, R., Holzschuh, A., Klein, A., Kleijn, D., Kremen, C., Landis, D., Laurance, W. and Westphal, C. (2012). Landscape moderation of biodiversity patterns and processes-eight hypotheses. Biological reviews of the Cambridge Philosophical Society 87: 661– 85, doi:10.1111/j.1469-185X.2011.00216.x.

Virtanen, P., Gommers, R., Oliphant, T. E., Haberland, M., Reddy, T., Cournapeau, D., Burovski, E., Peterson, P., Weckesser, W., Bright, J., van der Walt, S. J., Brett, M., Wilson, J., Millman, K. J., Mayorov, N., Nelson, A. R. J., Jones, E., Kern, R., Larson, E., Carey, C. J., Polat, İ., Feng, Y., Moore, E. W., VanderPlas, J., Laxalde, D., Perktold, J., Cimrman, R., Henriksen, I., Quintero, E. A., Harris, C. R., Archibald, A. M., Ribeiro, A. H., Pedregosa, F., van Mulbregt, P. and SciPy 1.0 Contributors (2020). SciPy 1.0: Fundamental Algorithms for Scientific Computing in Python. Nature Methods 17: 261–272, doi:10.1038/s41592-019-0686-2.

Walther, G.-R., Post, E., Convey, P., Menzel, A., Parmesan, C., Beebee, T., Fromentin, J.-M., Hoegh-Guldberg, O. and Bairlein, F. (2002). Ecological responses to recent climate change. Nature 416: 389–95, doi:10.1038/416389a.

Worboys, G., Francis, W. and Lockwood, M. (2010). Connectivity conservation management: A global guide, Connectivity Conservation Management: A Global Guide.. doi:10.4324/9781849774727.

Xue, J. and Su, B. (2017). Significant remote sensing vegetation indices: A review of developments and applications. Review Article — Open Access 2017: Article ID 1353691, doi:10.1155/2017/1353691.

Zha, Y., Gao, J. and Ni, S. (2003). Use of normalized difference built-up index in automatically mapping urban areas from tm imagery. International Journal of Remote Sensing - INT J REMOTE SENS 24: 583–594, doi:10.1080/01431160304987.

Zhao, Z., Zhang, S. hua, Xu, Z. yong, Bellisario, K., Dai, N. hua, Omrani, H. and Pijanowski, B. C. (2017). Automated bird acoustic event detection and robust species classification. Ecological Informatics 39: 99–108, 10.1016/j.ecoinf.2017.04.003.

Zhou, Y., Zheng, H., Huang, X., Hao, S., Li, D. and Zhao, J. (2022). Graph neural networks: Taxonomy, advances, and trends. ACM Transactions on Intelligent Systems and Technology 13: 1–54, doi:10.1145/3495161.

